# Geometry theory of distribution shapes for autoregulatory gene circuits

**DOI:** 10.1101/2024.04.02.587730

**Authors:** Ying Sheng, Genghong Lin, Feng Jiao, Chen Jia

## Abstract

In this study, we provide a complete mathematical characterization of the phase diagram of distribution shapes in an extension of the two-state telegraph model of stochastic gene expression in the presence of positive or negative autoregulation. Using the techniques of second-order difference equations and nonlinear discrete dynamical systems, we prove that the feedback loop can only produce three shapes of steady-state protein distributions (decaying, bell-shaped, and bimodal), corresponding to three distinct parameter regions in the phase diagram. The boundaries of the three regions are characterized by two continuous curves, which can be constructed geometrically by the contour lines of a series of ratio operators. Based on the geometric structure of the phase diagram, we then provide some simple and verifiable sufficient and/or necessary conditions for the existence of the bimodal parameter region, as well as the conditions for the steady-state distribution to be decaying, bell-shaped, or bimodal. Finally, we also investigate how the phase diagram is affected by the strength of positive or negative feedback.

## 1 Introduction

In living cells, the expression of a gene is often related to that of other genes, forming various gene regulatory networks [1]. One of the most common network motifs is an autoregulatory genetic feedback loop whereby protein expressed from a gene activates or represses its own transcription [2]. It has been estimated that 40% of all transcription factors in *Escherichia coli* self-regulate [3] with most of them participating in negative autoregulation [2]. Feedback regulation was also shown to significantly affect intrinsic gene expression noise [4–7] as well as the response and relaxation times of transcription networks [3, 8].

Over the past two decades, significant progress has been made in the mathematical modelling and analytical theory of autoregulated gene expression. Within the continuous-time Markovian framework, some studies model the protein dynamics explicitly and model the gene dynamics implicity [9–11]. In this case, both the steady-state [9–11] and time-dependent [12, 13] distributions of protein numbers can be solved exactly. Other studies have considered more realistic models where both the gene and protein dynamics are modelled explicitly [14–20]. Among these models, some assume that protein molecules are produced one at a time [14, 19], while others assume that protein molecules are produced in random bursts [15, 20]. Another distinguishing feature is that some models assume that there is no change in the protein number when a protein copy binds to a gene or when it unbinds [14, 15], while other models assume that the protein number decreases by one when a protein copy binds to a gene and increases by one when unbinding occurs [19, 20]. In the former case (binding fluctuations are igonored), the chemical master equation (𝒞ME) describing the autoregulatory loop can be solved analytically both in steady state [14–18] and in time [21, 22]. However, in the latter case (binding fluctuations are considered), only the steady-state solution has been obtained [19, 20] and the time-dependent solution is only obtained in the special case of fast gene state switching [13].

Experimentally, there are three commonly observed shapes for the steady-state distribution of protein numbers: a unimodal distribution with a zero peak (decaying distribution), a unimodal distribution with a nonzero peak (bell-shaped distribution), and a bimodal distribution with two peaks [23]. While the analytical theory for autoregulatory loops has been well developed, the exact protein distribution often involves various special functions. If protein expression occurs one at a time, then the exact solution is known to be a confluent hypergeometric function [14, 19]; if protein expression is bursty, then the exact solution is known to be a Gaussian hypergeometric function [15, 20]. In most cases, the shape of the protein distribution can only be determined by simulations due to the lack of mathematical results on the monotonicity of hypergeometric functions. A recent work [24] gave a rigorous classification of the three distribution shapes for the two-state telegraph model of stochastic gene expression; however, the authors did not take feedback regulation into account. Thus far, there is still a lack of a mathematical characterization of distribution shapes for autoregulatory gene circuits.

In this paper, we address this problem and provide a complete mathematical characterization of the phase diagram of distribution shapes in a minimal coupled positive-plus-negative feedback loop with gene state switching, protein synthesis, and protein degradation. The model includes the classical autoregulatory loops as special cases. In Sec. 2, the reaction scheme and the CME describing the stochastic dynamics of the feedback loop are introduced. In Sec. 3, we transform the CME into a second-order difference equation satisfied by the steady-state protein distribution and prove that the coupled gene circuit can only produce three distribution shapes (decaying, bell-shaped, and bimodal), which correspond to three distinct regions in the phase diagram. In Sec. 4, we introduce a series of ratio operators and obtain the nonlinear dynamical evolution satisfied by these operators. Crucially, we show that the boundaries of the three regions are characterized by two continuous increasing curves in the phase diagram, which can be constructed geometrically by the contour lines of these operators. Based on the geometric structure of the phase diagram, we also provide some simple and verifiable sufficient and/or necessary conditions for the existence of the bimodal parameter region, as well as the conditions for the steady-state distribution to be decaying, bell-shaped, or bimodal. In Sec. 5, we examine how the phase diagram is affected by the strengths of positive and negative feedback. We conclude in Sec. 6.

## 2 Model

Here we consider stochastic gene expression dynamics in a minimal coupled positive-plus-negative feedback loop with gene state switching, protein synthesis, and protein decay (Fig. 1(a)). Let *G* and *G*^*\**^ denote the inactive and active states of the gene and let *P* denote the corresponding protein. The protein is produced only when the gene is in the active state *G*^*\**^. Let *n* denote the copy number of protein. The reaction scheme underlying the coupled feedback loop is as follows: [16]:

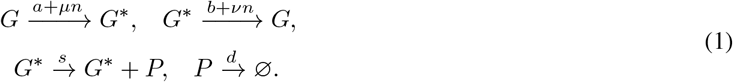

**Figure 1.**
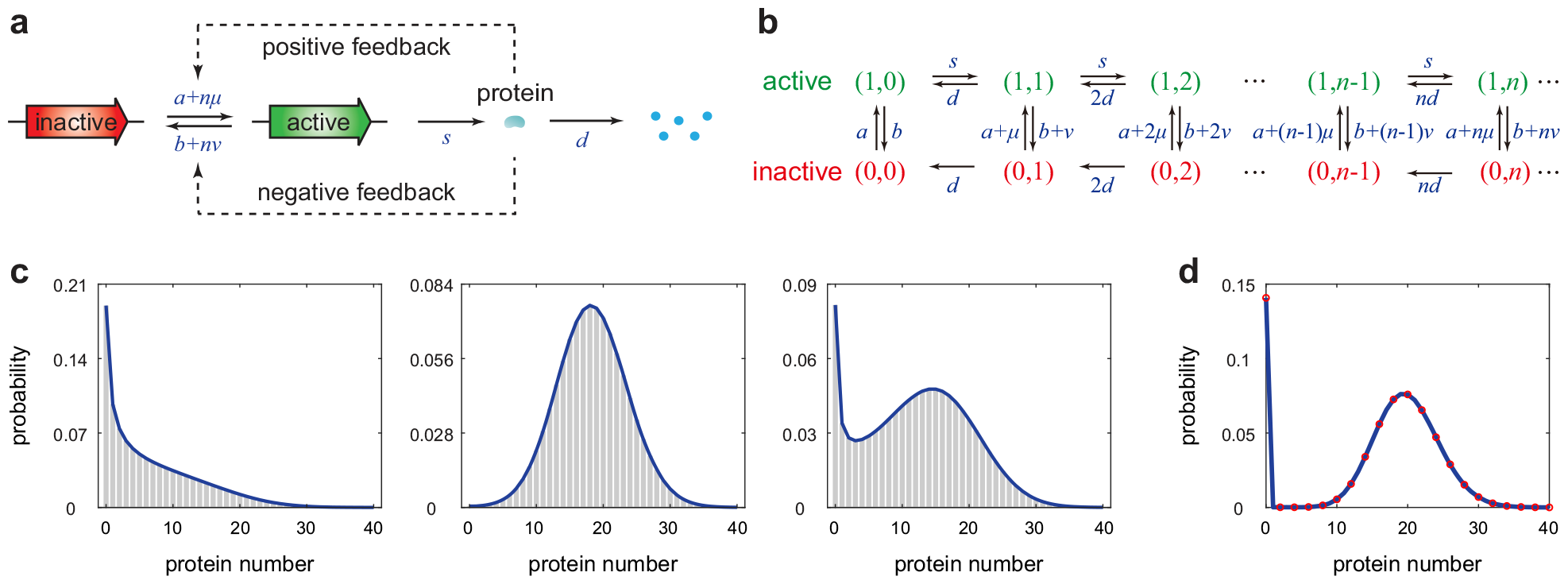
Model and its steady-state protein distributions. **(a)** A minimal coupled gene circuit with positive-plus-negative feedback. Protein molecules are produced only when the gene is active. **(b)** Transition diagram of the Markovian dynamics for the model illustrated in (a). **(c)** The model can generate three different shapes of steady-state protein distributions: a unimodal distribution with a zero peak (left panel), a unimodal distribution with a nonzero peak (middle panel), and a bimodal distribution with both a zero and a nonzero peak (right panel). The parameters are chosen as *s* = 30, *d* = 1, *a* = 0.5, *b* = 0.4, *µ* = 0, *ν* = 0.1 for the left panel, *s* = 30, *d* = 1, *a* = 1, *b* = 0.1, *µ* = 0.8, *ν* = 0.5 for the middle panel, and *s* = 30, *d* = 1, *a* = 0.4, *b* = 0.1, *µ* = 0.2, *ν* = 0.2 for the right panel. The distributions are computed using the finite-state projection (FSP) algorithm [28]. **(d)** When *a, b, ν «* 1, the protein number approximately has a zero-inflated Poisson (ZIP) distribution. The blue curve shows the simulated distribution obtained using FSP and the red circles show the approximated ZIP distribution given in Eq. (25). The parameters are chosen as *s* = 20, *d* = 1, *u* = 0.3, *a* = *b* = *ν* = 0.001.

Due to feedback regulation, the protein number *n* will directly or indirectly affect the switching rates between the two gene states. Here *a* and *b* are the spontaneous switching rates, *µ* and *ν* characterize the strengths of positive and negative feedback loops, respectively, *s* is the synthesis rate of protein in the active gene state, and *d* is the decay rate of protein either due to protein degradation or due to dilution during cell division [25–27]. Specifically, the protein decay rate can be represented as *d* = log 2*/T*_*p*_ + log 2*/T*_*c*_, where *T*_*p*_ is the protein half-life and *T*_*c*_ is the cell cycle duration. In what follows, for convenience, we set *d* = 1. This is not an arbitrary choice but stems from the fact that the time can always be re-normalized by the decay rate *d*. Specifically, the time *t* given below should be understood to be non-dimensional and equal to the real time multiplied by *d*, while the other parameters *s, a, b, µ*, and *ν* given below should also be understood to be non-dimensional and equal to their real values divided by *d*.

Note that the model considered here is more general than the classical model of autoregulatory feedback loops proposed by Hornos et al. [14]. In particular, the model describes a positive autoregulatory loop if the negative feedback strength *ν* vanishes, and it describes a negative autoregulatory loop if the positive feedback strength *µ* vanishes. In a coupled feedback loop, both *µ* and *ν* are nonzero. Coupled gene circuits widely exist in nature and they have been shown to be a crucial network motif to produce robust and tunable biological oscillations [29]. Biological examples of coupled feedback loops can be found in [16, 29].

The microstate of the gene can be represented by an ordered pair (*i, n*), where *i* is the gene state with *i* = 0, 1 corresponding to the inactive and active states, respectively, and *n* is the protein number. Let *P*_*i,n*_(*t*) denote the probability of having *n* protein molecules in an individual cell when the gene is in state *i*. Then the stochastic gene expression dynamics can be described by the Markov jump process shown in Fig. 1(b). The evolution of the Markovian dynamics is governed by the CMEs:

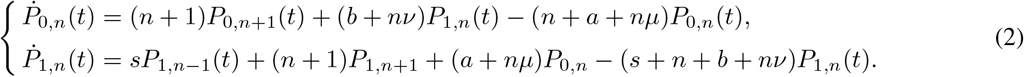

To proceed, let *P*_*n*_(*t*) = *P*_0,*n*_(*t*) + *P*_1,*n*_(*t*) denote the probability of having *n* protein molecules. For clarity, in what follows, we use *P*_*n*_(*t*) to represent the time-dependent protein distribution and use *P*_*n*_ to represent the steady-state protein distribution, i.e. *P*_*n*_ = lim_*t→∞*_ *P*_*n*_(*t*). In steady state, the CMEs given in Eq. (2) can be solved exactly and the steady-state distribution of protein numbers is given by [16, 18]

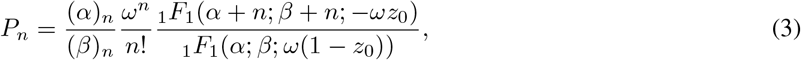

where (*x*)_*n*_ = *x*(*x* + 1) *· · ·* (*x* + *n −* 1) denotes the Pochhammer symbol, _1_*F*_1_(*α*; *β*; *z*) is Kummer’s confluent hypergeometric function, and

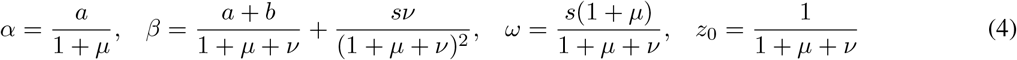

are four constants. Note that when *ν* = 0 or *µ* = 0, the coupled gene circuit reduces to an autoregulatory feedback loop, and the analytical distribution given in Eq. (3) reduces to the one obtained in [14].

## 3 Distribution shapes for coupled gene circuits

In experiments, there are three commonly observed shapes for the steady-state protein distribution: a unimodal distribution with a zero peak (decaying distribution), a unimodal distribution with a nonzero peak (bell-shaped distribution), and a bimodal distribution with both a zero and a nonzero peak [23]. According to simulations, our model is capable of producing all these three distribution shapes (Fig. 1(c)). Among the three shapes, the bimodal one is of particular interest since it separates isogenic cells into two distinct phenotypes [30, 31]. Mathematically, an important question is whether it is possible for a coupled gene circuit to produce other shapes of steady-state distributions such as a bimodal distribution with two nonzero peaks and a trimodal distribution [32]. To answer this question, a direct analysis of the analytical distribution given in Eq. (3) is extremely difficult since (i) mathematical results on the monotonicity of the confluence hypergeometric function are very limited and (ii) the dependence of the constants *α, β, ω*, and *z*_0_ on the parameters *s, a, b, µ*, and *ν* are highly nonlinear. To solve this problem, we transform the CMEs into a recurrence relation between *P*_*n−*1_, *P*_*n*_, and *P*_*n*+1_, as shown in the following lemma.

### Lemma 3.1

The stationary distribution *P*_*n*_ satisfies the following recurrence relation:

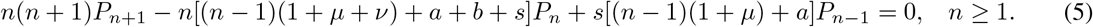

*Proof*. We introduce the generating functions

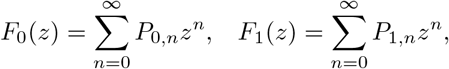

and let *F* (*z*) = *F*_0_(*z*) + *F*_1_(*z*). In steady state, the CMEs given in Eq. (2) can be converted into the following set of first-order linear ordinary differential equations (ODEs) satisfied by the generating functions [16]:

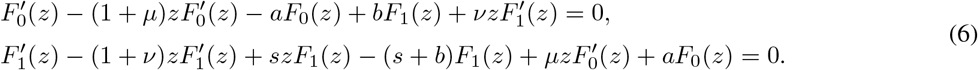

Adding the two identities in Eq. (6), we obtain *F* ′(*z*) = *sF*_1_(*z*). Inserting this into the second identity in Eq. (6) and applying *F*_0_(*z*) = *F* (*z*) *− F*_1_(*z*), we find that *F* (*z*) satisfies the following second-order ODE:

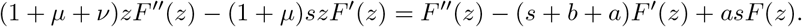

Taking high-order derivatives on both sides of the above equation, it can be proved by induction that *F* ^(*n*)^(*z*) satisfies the following recurrence relation:

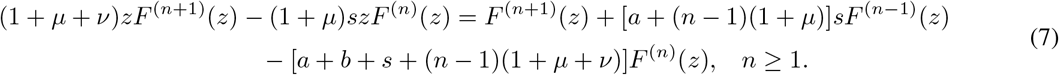

Recall that the steady-state distribution can be recovered by taking derivatives of the generating function at *z* = 0, i.e.

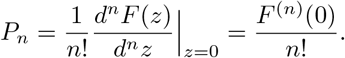

The desired result then follows from Eq. (7) by taking *z* = 0.

For clarity, we now give the rigorous definitions of the three distribution shapes. Mathematically, the steady-state distribution *P*_*n*_ is said to be *decaying* if

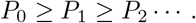

It is said to be *bell-shaped* if there exists *n*_1_ *≥* 1 such that

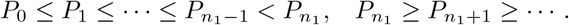

It is said to be *bimodal* with a zero and a nonzero peak if there exist *n*_1_ *> n*_0_ *≥* 1 such that

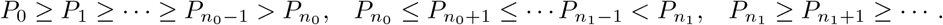

Next we analyze the shape of the steady-state distribution using the technique of difference equations. Note that by introducing the difference operator Δ*P*_*n*_ = *P*_*n*_ *− P*_*n−*1_, the recurrence relation given in Eq. (5) can be rewritten as the following non-autonomous second-order difference equation:

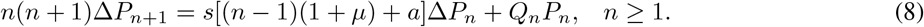

where

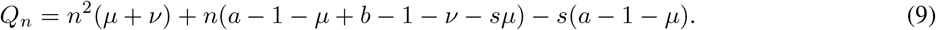

The following theorem shows that a coupled gene circuit can only produce the above three distribution shapes.

### Theorem 3.2

In steady state, the coupled gene circuit described by Eq. (1) can only produce a decaying distribution, a bell-shaped distribution, or a bimodal distribution with a zero and a nonzero peak (Fig. 1(c)).

*Proof*. To prove the desired result, we only need to show that *P*_*n*_ peaks twice at most, with possible modes at *n* = 0 and *n* = *n*_1_ *>* 0. Note that *n ≥* 1 is a nonzero mode of *P*_*n*_ if and only if Δ*P*_*n*_ *>* 0 and Δ*P*_*n*+1_ *≤* 0. Let *n*_1_ be the smallest nonzero mode of *P*_*n*_. Suppose that *P*_*n*_ has another nonzero mode *n*_2_. Clearly, we have *n*_2_ *≥ n*_1_ + 2 and

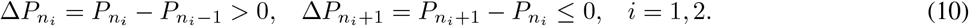

Combining these inequalities with Eq. (8), we find that 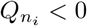 for *i* = 1, 2. From Eq. (9), it is easy to see that *Q*_*n*_ is convex with respect to *n*. This suggests that *Q*_*n*_ *<* 0 for all *n* ∈ [*n*_1_, *n*_2_]. Since *n*_1_ and *n*_2_ are two modes of *P*_*n*_, there must exist *m* ∈ (*n*_1_, *n*_2_) such that *P*_*n*_ achieves its minimum at *n* = *m*, i.e.

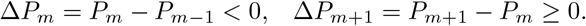

Inserting these inequalities into Eq. (8), we find that *Q*_*m*_ *≥* 0. This contradicts our previous observation that *Q*_*n*_ *<* 0 for all *n* ∈ [*n*_1_, *n*_2_]. This indicates that *P*_*n*_ cannot have two nonzero modes. In other words, it peaks twice at most, with possible modes at *n* = 0 and *n* = *n*_1_ *>* 0. This completes the proof of the theorem.

## 4 Phase diagrams for coupled gene circuits

To analyze how model parameters influence the shape of the steady-state distribution *P*_*n*_, we introduce the ratio

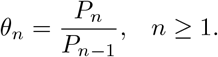

The three distribution shapes can also be analyzed using *θ*_*n*_. For example, if 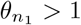 and 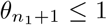, then *P*_*n*_ has a mode at *n* = *n*_1_. To analyze *θ*_*n*_, we first divide Eq. (8) by *P*_*n*_ and obtain

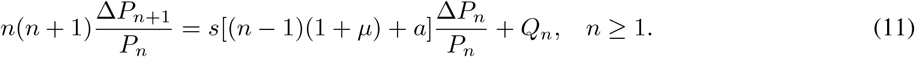

Note that Δ*P*_*n*+1_*/P*_*n*_ = *θ*_*n*+1_ *−* 1 and Δ*P*_*n*_*/P*_*n*_ = 1 *−* 1*/θ*_*n*_. Inserting these into Eq. (11), we find that *θ*_*n*_ satisfies the following nonlinear discrete dynamical system:

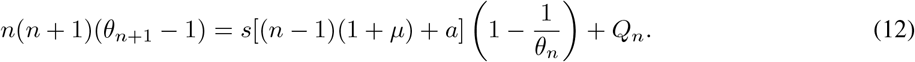

For convenience, we rewrite *θ*_*n*_ as *θ*_*n*_(*a, b*) to emphasize its dependence on *a* and *b*. It then follows from Eq. (3) that

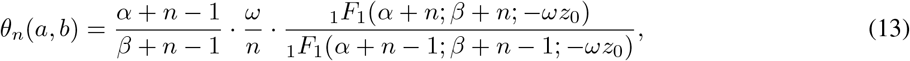

where

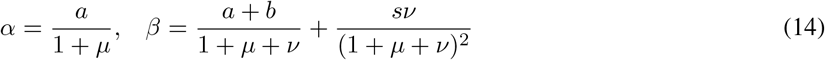

are the constants defined in Eq. (4). Note that if we replace *a* by *a* + 1 + *µ* and replace *b* by *b* + *ν* in Eq. (14), then *α* is replaced by *α* + 1 and *β* is replaced by *β* + 1. This clearly shows that

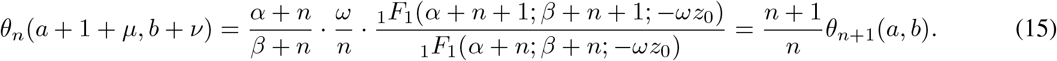

Finally, inserting Eq. (15) into Eq. (12) yields

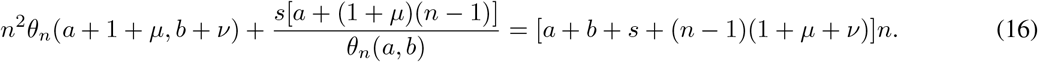

These equations will play a crucial role in analyzing the properties of *θ*_*n*_(*a, b*) (see Lemma 4.3 below and the lemmas in Appendix).

### Lemma 4.1

For any *s, a, b >* 0 and *µ, ν ≥* 0, we have

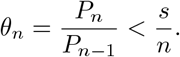

*Proof*. The proof of this lemma is complicated and can be found in Supplementary Section 1.

The above lemma shows that *P*_*n*_ *< P*_*n−*1_ for all *n ≥ s*. Hence we only need to analyze the monotonicity of *P*_*n*_ within the interval *n* ∈ [0, *s*). This implies the following theorem.

### Theorem 4.2

If *s ≤* 1, then the distribution *P*_*n*_ must be decaying.

*Proof*. If *s ≤* 1, then *P*_*n*_ *< P*_*n−*1_ for all *n ≥* 1, according to Lemma 4.1. This gives the desired result.

Recall that we have assumed that *d* = 1. Hence *s* = *s/d* represents the typical protein number in the active gene state. The condition *s ≤* 1 is rare since the protein number in a single cell is typically larger than 1. We next consider the more common case of *s >* 1. To this end, we first prove the strict monotonicity of *θ*_*n*_ with respect to the spontaneous switching rates *a* and *b*, as shown in the following lemma.

### Lemma 4.3

Let *µ, ν ≥* 0 and *s >* 0 be fixed. For any *n ≥* 1, *θ*_*n*_(*a, b*) strictly increases with respect to *a* and strictly decreases with respect to *b*.

*Proof*. For convenience, set

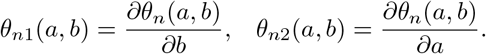

Differentiating Eq. (16) with respect to *b*, we obtain

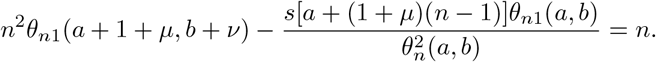

This clearly shows that

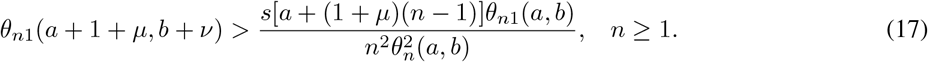

Moreover, differentiating Eq. (16) with respect to *a* yields

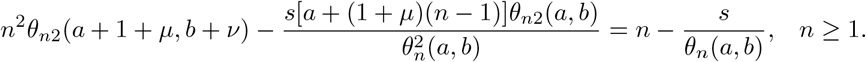

### Lemma 4.1

shows that the right-hand side of the above equation must be negative. Hence we obtain

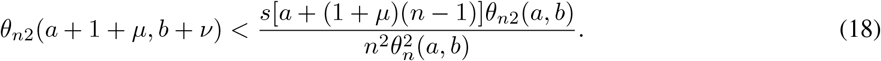

We now prove that *θ*_*n*1_(*a, b*) *<* 0 for any *a, b >* 0 and *n ≥* 1. If this does not hold, then there exist *a*_0_, *b*_0_ *>* 0 such that *θ*_*n*1_(*a*_0_, *b*_0_) *≥* 0. Applying Eq. (17) repeatedly, we have *θ*_*n*1_(*a*_*k*_, *b*_*k*_) *>* 0 for any *k ≥* 1, where *a*_*k*_ = *a*_0_ + *k*(1 + *µ*) and *b*_*k*_ = *b*_0_ + *kν*. Take a sufficiently large *K ≥* 1 and a sufficiently small *ϵ >* 0 such that

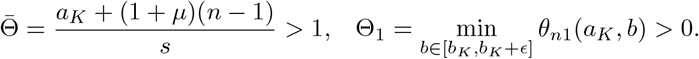

Replacing *a* and *b* by *a*_*K*_ + (1 + *µ*)(*k −* 1) and *b* + *ν*(*k −* 1), respectively, in Eq. (17), we obtain

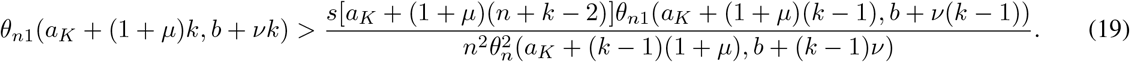

For any *b ∈* [*b*_*K*_, *b*_*K*_ + *ϵ*], since *θ*_*n*1_(*a*_*K*_, *b*) *<* 0, it follows from Eq. (17) that *θ*_*n*1_(*a*_*K*_ + (1 + *µ*)*k, b* + *νk*) *>* 0. In addition, since *θ*_*n*_ *< s/n*, it follows from Eq. (19) that

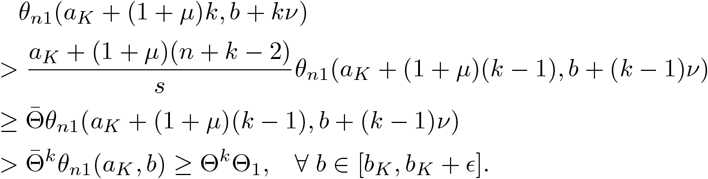

By the mean value theorem, there exists *b* ∈ (*b*_*K*_, *b*_*K*_ + *ϵ*) such that

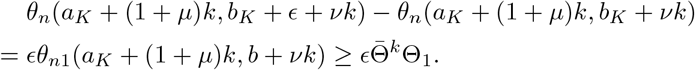

Since *θ*_*n*_ *< s/n*, the left-hand side of the above equation is uniformly bounded for all *k ≥* 1, while the right-hand side tends to infinity as *k → ∞*. This yields a contradiction and hence *θ*_*n*1_(*a, b*) *<* 0 for all *a, b >* 0 and *n ≥* 1. This shows that *θ*_*n*_(*a, b*) is strictly decreasing with respect to *b*.

Similarly, using Eq. (18), it can be proved that *θ*_*n*2_(*a, b*) *>* 0 for any *a, b >* 0 and *n ≥* 1 and hence *θ*_*n*_(*a, b*) is strictly increasing with respect to *a*.

We next examine the phase diagram of distribution shapes. Note that for any *n ≥* 1, the equation *θ*_*n*_(*a, b*) = 1 determines a contour line of *θ*_*n*_ in the *b*-*a* plane. The strict monotonicity of *θ*_*n*_(*a, b*) with respect to *a* and *b* guarantees that the contour line *θ*_*n*_(*a, b*) = 1 is a smooth, strictly increasing curve in the *b*-*a* plane (this follows directly from the implicit function theorem [33]). We make a crucial observation that if we tune the parameters *s, µ*, and *ν*, then the steady-state distribution *P*_*n*_ may change from one shape to another. In this case, value of *θ*_*n*_ must cross the contour line *θ*_*n*_(*a, b*) = 1 for some *n ≥* 1. Hence we can use the contour lines of *θ*_*n*_ to determine the boundaries of the three distribution shapes in the *b*-*a phase plane*. Roughly speaking, we will use the contour lines of *θ*_*n*_ to construct two continuous curves 𝒞_1_ and 𝒞_2_, which separate the phase plane into three regions, corresponding to the three distinct distribution shapes. This forms a *phase diagram* for coupled gene circuits, as illustrated in Fig. 2(b),(c).

**Figure 2.**
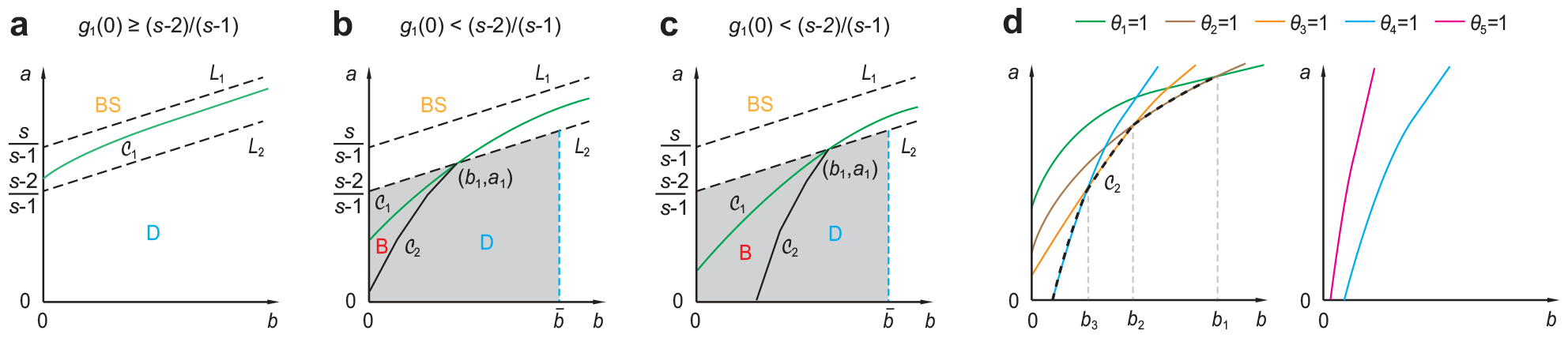
Phase diagrams for coupled gene circuits in the *b*-*a* plane. Here *a* and *b* are the spontaneously switching rates between the two gene states. When *s >* 1, the curve *𝒞*_1_ exists and lies between the straight lines *L*_1_ and *L*_2_. **(a)** When *g*_1_(0) *≥* (*s−*2)*/*(*s−*1), the curve *𝒞*_1_ separates the phase diagram into a decaying (D) and a bell-shaped (BS) region. **(b),(c)** When *g*_1_(0) *<* (*s −* 2)*/*(*s −* 1), the curve *𝒞*_2_ also exists, and the two curves *𝒞*_1_ and *𝒞*_2_ separate the phase diagram into a decaying (D), a bell-shaped (BS), and a bimodal (B) region. The curve *𝒞*_2_ intersects with *𝒞*_1_ and *L*_2_ at a unique point (*b, a*) = (*b*_1_, *a*_1_). It may start from the *a*-axis, as illustrated in (b), or start from the *b*-axis, as illustrated in (c). **(d)** Construction of the curves *𝒞*_1_ and *𝒞*_2_. The curve *𝒞*_1_ is the contour line of *θ*_1_ (green curve). In Appendix, we prove that there exists an integer 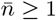 (the figure shows the case of 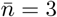) such that for each 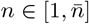, the contour lines of *θ*_*n*_ and *θ*_*n*+1_ intersect at a unique point (*b*_*n*_, *a*_*n*_) in the *b*-*a* plane (left panel), while the contour lines of 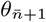 and 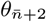 do not intersect (right panel). Then the curve *𝒞*_2_ (black dashed curve) is consisted of (i) the truncated contour lines of *θ*_*n*_ between (*b*_*n−*1_, *a*_*n−*1_) and (*b*_*n*_, *a*_*n*_) for all 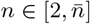 and (ii) the truncated contour line of 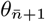 between 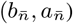 and its intersection point with the coordinate axes.

Specifically, we have the following theorem, which characterizes the geometric structure of the phase diagram for any *s >* 1. This is also the main result of the present paper.

### Theorem 4.4

Let *s >* 1 and *µ, ν ≥* 0 be fixed and let *ω* be the constant given in Eq. (4). Then *θ*_1_(*a, b*) = 1 determines a smooth and strictly increasing contour line 𝒞_1_ : *a* = *g*_1_(*b*), *b >* 0 in the *b*-*a* plane, which satisfies

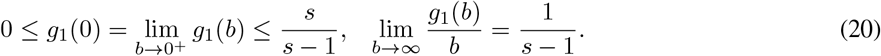

In particular, *g*_1_(0) = 0 if and only if *ν* = 0, and

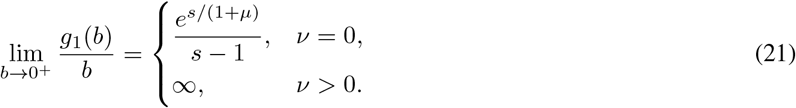

Specifically, we have the following two cases.

i. If *g*_1_(0) *≥* (*s −* 2)*/*(*s −* 1), then 𝒞_1_ lies between the two straight lines

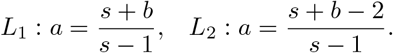

The distribution *P*_*n*_ is bell-shaped in the parameter region above 𝒞_1_, and the distribution is decaying in the parameter region below 𝒞_1_ (see Fig. 2(a) for an illustration).
ii. If *g*_1_(0) *<* (*s −* 2)*/*(*s −* 1), then *ω >* 2 and 𝒞_1_ intersects with *L*_2_ at a unique point (*b, a*) = (*b*_1_, *a*_1_) with

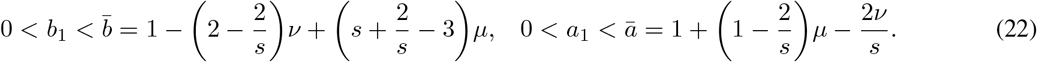

In particular, *𝒞*_1_ lies below *L*_2_ for *b < b*_1_ and lies between *L*_1_ and *L*_2_ for *b > b*_1_. There also exists another piecewise smooth and strictly increasing curve *𝒞*_2_ : *a* = *g*(*b*), *b*^*\**^ *< b ≤ b*_1_ in the *b*-*a* plane with *b*^*\**^ = 0 (see Fig. 2(b) for an illustration) or *b*^*\**^ *>* 0 (see Fig. 2(c) for an illustration). Specifically, *𝒞*_2_ is constructed by splicing 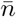 contour lines (this will be explained in later in Remark 4.5), i.e. the contour lines *θ*_*n*_(*a, b*) = 1 for 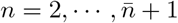, where 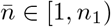 is an integer with

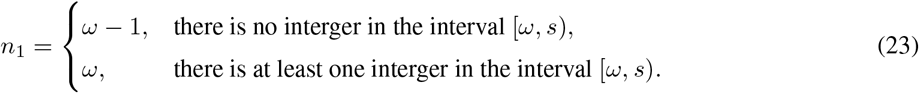

In particular, *𝒞*_2_ intersects with *𝒞*_1_ and *L*_2_ at a unique point (*b, a*) = (*b*_1_, *a*_1_) and *𝒞*_2_ lies below *𝒞*_1_ for *b < b*_1_. The distribution *P*_*n*_ is bell-shaped in the parameter region above *𝒞*_1_; the distribution is decaying in the parameter region below *𝒞*_1_ and on the right-hand side of *𝒞*_2_; the distribution is bimodal in the parameter region enclosed by *𝒞*_1_, *𝒞*_2_, and the coordinate axes (Fig. 2(b),(c)).

*Proof*. The proof of this theorem requires many lemmas and is very complicated. These statements of these lemmas and the proof of the theorem can be found in Appendix.

### Remark 4.5.

We emphasize that in Fig. 2(b),(c), the curves *𝒞*_1_ and *𝒞*_2_ separating the three phases (distribution shapes) intersect at a unique point (*b, a*) = (*b*_1_, *a*_1_); this is analogous to the triple point in physics and chemistry where all three phases of a substance (gas, liquid, and solid) coexist in thermodynamic equilibrium [34]. The curve *𝒞*_1_ is simply the contour line of *θ*_1_. The construction of *𝒞*_2_ is much more complicated and is intuitively shown in Fig. 2(d). In Appendix, we prove that there exists an integer 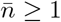 such that for each 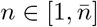, the contour lines of *θ*_*n*_ and *θ*_*n*+1_ intersect at a unique point (*b*_*n*_, *a*_*n*_) (left panel), while the contour lines of 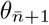 and 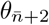 do not intersect (right panel). Then the curve *𝒞*_2_ (shown by the black dashed curve) is consisted of (i) the truncated contour lines of *θ*_*n*_ between (*b*_*n−*1_, *a*_*n−*1_) and (*b*_*n*_, *a*_*n*_) for all 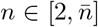 and (ii) the truncated contour line of 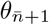 between 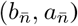 and its intersection point with the coordinate axes.

In summary, Theorem 4.2 and 4.4 give a complete geometric characterization of the phase diagram for coupled gene circuits. When *s ≤* 1, Theorem 4.2 shows that the steady-state distribution is always decaying. When *s >* 1, all the three distribution shapes may occur and Theorem 4.4 gives the exact boundaries between them. Specifically, the curve *𝒞*_1_ separates the phase diagram into one region for bell-shaped distributions (bell-shaped region) and the other region for decaying and bimodal distributions. The curve *𝒞*_2_, if it exists, further separates the latter region into one subregion for decaying distributions (decaying region) and the other subregion for bimodal distributions (bimodal region). Theorem 4.4 also gives the sufficient and necessary condition for the existence of the bimodal region — it exists if and only of *g*_1_(0) *<* (*s −* 2)*/*(*s −* 1). However, it is very difficult to verify this condition since *g*_1_(0) in general cannot be computed explicitly. The following theorems gives a simple sufficient and necessary condition that guarantees the existence of the bimodal region for positive feedback loops.

### Theorem 4.6

Let *µ ≥* 0 be fixed. For positive feedback loops (*ν* = 0), the bimodal region exists in the *b*-*a* plane if and only if *s >* 2.

*Proof*. Theorem 4.2 shows that there is no bimodal region when *s ≤* 1. When *s >* 1, Theorem 4.4 shows that the bimodal region exists if and only if *g*_1_(0) *<* (*s −* 2)*/*(*s −* 1). By Theorem 4.4, *ν* = 0 if and only if *g*_1_(0) = 0. Hence when *ν* = 0, the bimodal region exists if and only if (*s −* 2)*/*(*s −* 1) *>* 0, i.e. *s >* 2.

The following theorem give the sufficient and necessary condition for bimodality when *a, b, ν* ≪ 1.

### Theorem 4.7

Let *s >* 2 and *µ ≥* 0 be fixed, and let *a, b, ν* ≪ 1 be sufficiently small. Then the distribution *P*_*n*_ is bimodal if and only if

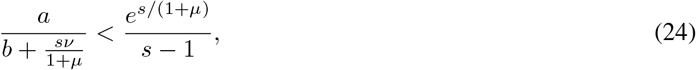

and it is bell-shaped otherwise.

*Proof*. When *a, b, ν* ≪ 1, using the standard two-time-scale simplification technique for Markov chains [35–37], it can be proved that the protein number approximately has a zero-inflated Poisson (ZIP) distribution [38, 39] (see Supplementary Section 2 for the proof and see Fig. 1(d) for the comparison between the real and approximated distributions)

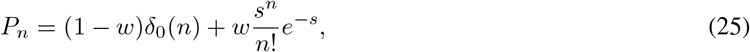

where *δ*_0_(*n*) is Kronecker’s delta which takes the value of 1 when *n* = 0 and takes the value of 0 otherwise, and

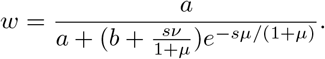

Note that this is a mixture of a point mass at zero and a Poisson distribution. Since a Poisson distribution is unimodal, the mixture of a point mass at zero and a Poisson distribution can yield a bimodal distribution (Fig. 1(d)). To see when the ZIP distribution is bimodal, note that the mode of the Poisson part is given by

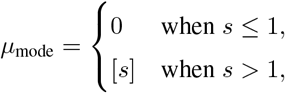

where ⌈*s⌉* denotes the smallest integer greater than *s*. Hence the ZIP distribution peaks at both zero and the non-zero mode ⌈*s⌉* if and only if *P*_0_ *> P*_1_ and *µ*_mode_ *>* 2, i.e.

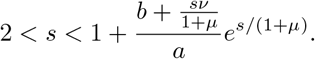

Otherwise, it is bell-shaped. This gives the desired result.

The following theorems gives a simple and verifiable *sufficient condition* for the existence of the bimodal region for general couple gene circuits.

### Theorem 4.8

Let *µ ≥* 0 be fixed. Then the bimodal region exists in the *b*-*a* plane if *s >* 2 and *ν ≪* 1.

*Proof*. When *s >* 2 and *ν ≪* 1, we can always find *a, b ≪* 1 such that Eq. (24) holds. Hence the desired result follows from Theorem 4.7.

The following theorem gives some simple and verifiable *necessary conditions* for the existence of the bimodal region for general couple gene circuits.

### Theorem 4.9

The bimodal region does not exist in the *b*-*a* plane if any one of the following three conditions is satisfied:

i. *s ≤* 2;
ii. *s >* 2 and (*s −* 2)(1 + *µ*) *−* 2*ν ≤* 0;
iii. *s >* 2 and (*s −* 2)*µ −* 2*ν* + *s/*(*s −* 1) *≤* 0.

*Proof*. Theorem 4.2 shows that the bimodal region does not exist when *s ≤* 1. When *s >* 1, Theorem 4.4 shows that the bimodal region exists if and only if *g*_1_(0) *<* (*s −* 2)*/*(*s −* 1). When 1 *< s ≤* 2, we have *g*_1_(0) *≥* 0 *≥* (*s −* 2)*/*(*s −* 1), and thus the bimodal region does not exist in the case. This proves (i).

Let *s >* 2 be fixed. Recall that the bimodal region exists in the *b*-*a* plane if and only if the curve *𝒞*_2_ exists. There are two cases where *𝒞*_2_ does not exist, according to Theorem 4.4. First, if *𝒞*_2_ exists, then we must have *ω >* 2. This is impossible if *ω ≤* 2, i.e.

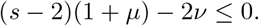

Second, if *𝒞*_2_ exists, then it must intersect with *𝒞*_1_ and *L*_2_ at 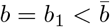 (Fig. 2(b),(c)). This is impossible if 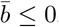, i.e.

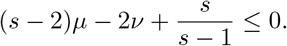

This proves (ii) and (iii).

When *s >* 2, the system may produce a bimodal steady-state distribution. If the gene is unregulated (*µ* = *ν* = 0), it has been proved in [24] that bimodality can only occur when both the gene switching rates are smaller than the degradation rate, i.e. *a, b <* 1. The following corollary generalizes this result to coupled gene circuits.

### Corollary 4.10.

Let *µ, ν ≥* 0 and *s >* 2 be fixed. Then a coupled feedback loop can exhibit bimodality only when

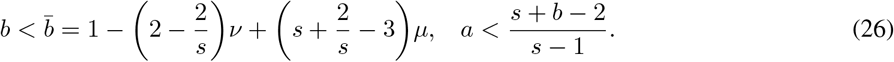

In particular, for negative feedback loops (*µ* = 0), bimodality can only occur when

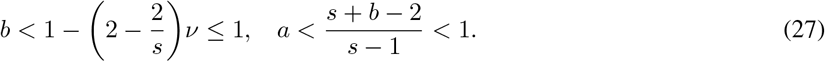

*Proof*. By Theorem 4.2, if *𝒞*_2_ exists, then the bimodal region is enclosed by *𝒞*_1_, *𝒞*_2_, and the coordinate axes. Since *𝒞*_1_ and *𝒞*_2_ lie below *L*_2_ for *b < b*_1_, the bimodal region must lie below the straight line *L*_2_. Hence in order for bimodality to occur, we must have

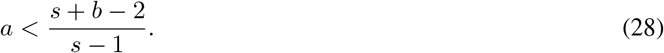

Furthermore, since *𝒞*_2_ is increasing and since 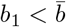, the bimodal region must lie on the left-hand side of the vertical line 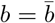. This gives the desired result.

The above corollary quantifies the “potential” bimodal region in the phase diagram, i.e. the region enclosed by the straight line *L*_2_ and the vertical line 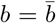 (see the grey region in Fig. 2(b),(c)). In other words, bimodality can never occur when the parameter set falls out of the potential bimodal region. In particular, a negative feedback loop can exhibit bimodality only when *a, b <* 1. The following corollary characterizes the parameter regions for decaying and bell-shaped distributions.

### Corollary 4.11.

Let *s >* 1 and *µ, ν ≥* 0 be fixed. Then the distribution *P*_*n*_ is decaying if

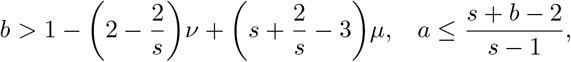

and the distribution is bell-shaped if

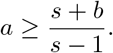

*Proof*. From Fig. 2(a)-(c), the distribution must be decaying in the region below the straight line *L*_2_ and on the right-hand side of the vertical line 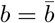. This proves the first part of the corollary. On the other hand, the distribution must be bell-shaped in the region above the straight line *L*_1_. This proves the second part of the corollary.

## 5 Further analysis on the phase diagram

It has long been known that positive feedback loops tend to promote bimodality, while negative feedback loops tend to restrain bimodality [40, 41]. Our theory provides a quantitative characterization of this fact. Corollary 4.10 shows that when *s >* 2, bimodality can occur only when the parameter set lies in the potential bimodal region enclosed by the straight line *L*_2_ and the vertical line 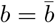, i.e.

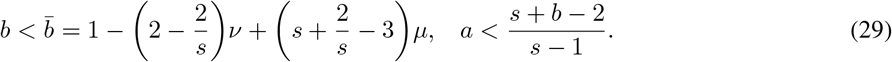

As the positive (negative) feedback strength *µ* (*ν*) increases, the straight line *L*_2_ remains invariant, while the vertical line 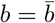 moves to the right (left) in the phase plane. Hence the intersection of *𝒞*_1_ and *𝒞*_2_ moves up (down) along *L*_2_, forcing the potential bimodal region to enlarge (shrink) (Fig. 2(b),(c)). This coincides with previous results that positive feedback is more likely to enlarge the bimodal region, while the negative feedback is more likely to shrink it. This can be also seen from Theorem 4.9, where we have shown that when *s >* 2, bimodality can never occur when

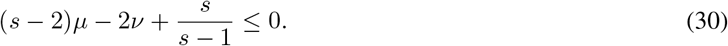

Clearly, as the negative feedback strength *ν* increases or as the positive feedback strength *µ* decreases, this condition is more easily to be satisfied. This again shows that bimodality is restrained by negative feedback and is promoted by positive feedback.

Fig. 3 compares the phase diagrams for positive, negative, and coupled feedback loops under different feedback strengths (weak, intermediately strong, and strong feedback). By Theorem 4.4, for positive feedback loops (*ν* = 0), we have *g*_1_(0) = 0 which means that the curve *𝒞*_1_ starts from the origin (Fig. 3(a)); for negative and coupled feedback loops (*ν >* 0), we have *g*_1_(0) *>* 0 which means that *𝒞*_1_ starts from the *a*-axis (Fig. 3(b),(c)). This is a crucial difference between positive and negative/coupled feedback loops. In addition, when *s >* 2, Theorem 4.6 show that the bimodal region always exists for positive feedback loops (Fig. 3(a)). For negative and coupled feedback loops, the bimodal region exists when the negative feedback is weak, according to Theorem 4.8. However, the bimodal region may disappear when negative feedback is sufficiently strong (Fig. 3(b)). This is another difference between positive and negative/coupled gene circuits.

**Figure 3.**
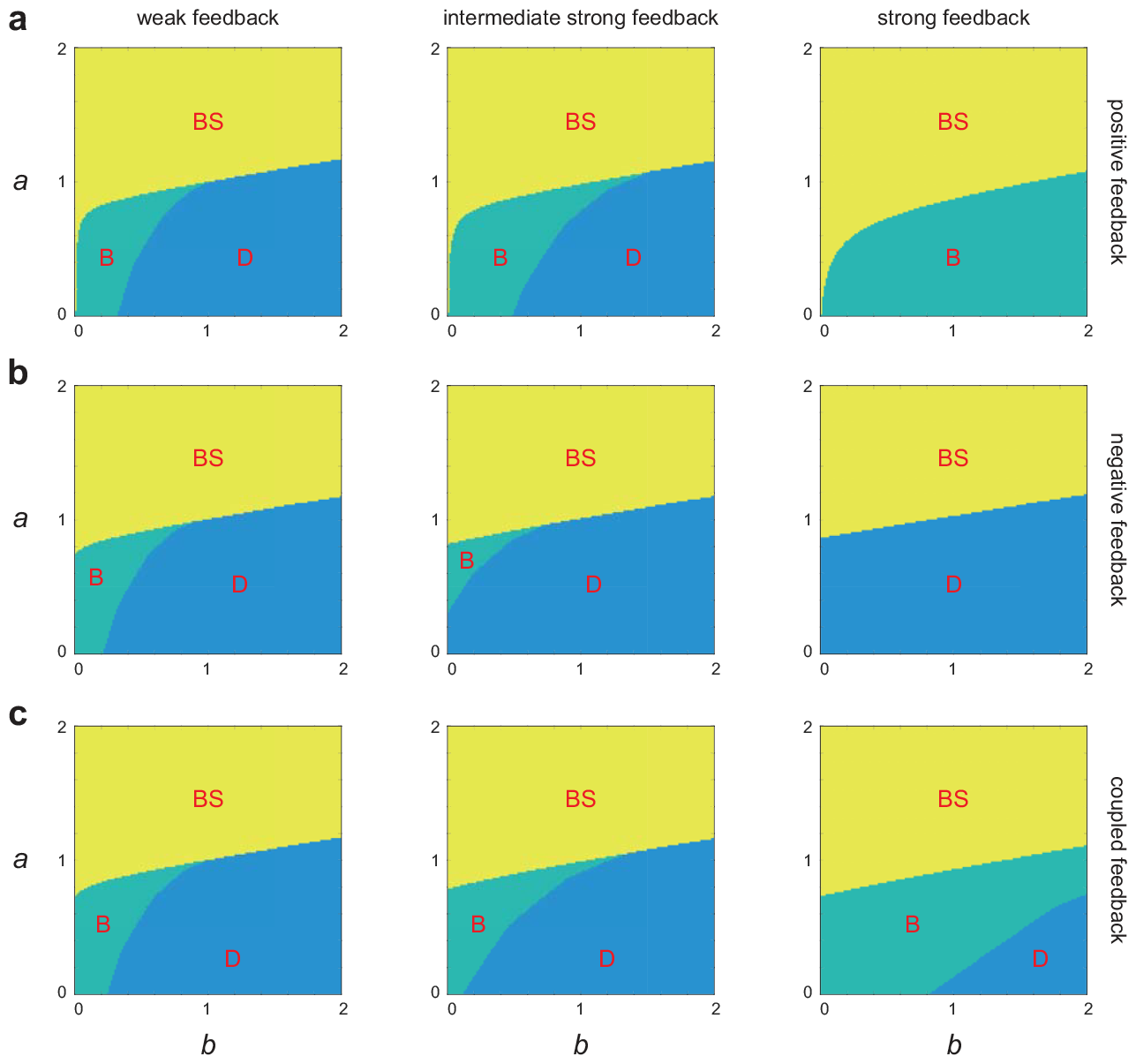
Phase diagrams in the *a*-*b* plane under different feedback controls and feedback strengths. The system can produce a decaying (D), a bell-shaped (BS), or a bimodal (D) steady-state protein distribution. **(a)** Positive feedback loops. The parameters are chosen as *s* = 8, *µ* = 0.0205 for weak feedback, *µ* = 0.123 for intermediate strong feedback, and *µ* = 0.738 for strong feedback. **(b)** Negative feedback loops. The parameters are chosen as *s* = 8, *ν* = 0.0115 for weak feedback, *ν* = 0.069 for intermediate strong feedback, and *ν* = 0.414 for strong feedback. **(c)** Coupled feedback loops. The parameters are chosen as *s* = 8, *µ* = 0.0205, *ν* = 0.0115 for weak feedback, *µ* = 0.123, *ν* = 0.414 for intermediate strong feedback, and *µ* = 0.738, *ν* = 0.069 for strong feedback.

We next quantitatively analyze how the bimodal region is affected by the feedback strength. By Theorem 4.4, for the positive feedback case (*ν* = 0), the slope of the curve *𝒞*_1_ at the origin is given by

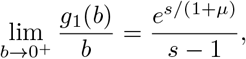

which decreases as the feedback strength *µ* increases. Hence as positive feedback becomes stronger, *𝒞*_1_ becomes flatter due to a smaller slope at the origin and *𝒞*_2_ moves to the right (Fig. 3(a)), forming larger bimodal and bell-shaped regions and a smaller decaying region. In particular, when *s, µ ≫ a, b, ν*, Theorem 4.7 shows that the system displays bimodality if and only if Eq. (24) holds.

For the negative feedback case, we have seen that *𝒞*_1_ always starts from the *a*-axis. When negative feedback is weak, *𝒞*_2_ starts from the *b*-axis and hence the bimodal region contains the origin (Fig. 3(b)). As the feedback strength *µ* increases, *𝒞*_1_ slightly moves up, and the starting point of *𝒞*_2_ crosses the origin and then moves to the *a*-axis. In this case, the bell-shaped region remains almost unchanged and the bimodal region converts to the decaying region. When negative feedback is sufficiently strong, the bimodal region disappears. When *s >* 2, it follows from (30) that the bimodal region disappears when

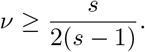

For the coupled feedback case, as the feedback strengthes *µ* and *ν* increase simultaneously, the bimodal region may either enlarge or shrink. When *s >* 2, it follows from Eq. (29) that when

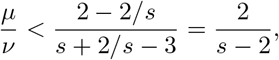

the vertical line 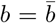 moves to the left in the phase plane. In this case, the potential bimodal region shrink and the actual bimodal region also becomes smaller. On the contrary, the bimodal region enlarges when *µ/ν >* 2*/*(*s −* 2). In particular, when *µ* and *ν* both increase while maintaining equal (*µ* = *ν*), the bimodal region shrinks when *s <* 4 and enlarges when *s >* 4 (Fig. 3(c)).

## 6 Conclusions and discussion

In the present paper, we provided a complete mathematical characterization of the phase diagram of distribution shapes in a minimal coupled positive-plus-negative feedback loop with gene state switching, protein synthesis, and protein decay [16]. This coupled gene circuit includes positive and negative autoregulatory feedback loops as special cases. The CMEs for this model can be solved exactly in steady state [14, 18] and the analytical solution is widely known to be a confluent hypergeometric function. However, a direct analysis of the exact protein distribution is extremely difficult since mathematical results on the monotonicity of the confluence hypergeometric function are quite limited. To overcome this difficulty, we transformed the CMEs into a second-order difference equation satisfied by the steady-state distribution *P*_*n*_ and a nonlinear discrete dynamical system satisfied by the ratio *θ*_*n*_ = *P*_*n*_*/P*_*n−*1_. Using the techniques of difference equations, we rigorously prove that the coupled gene circuit can only produce three distinct distribution shapes (decaying, bell-shaped distribution, and bimodal).

Furthermore, an in-depth and intricate analysis of the nonlinear discrete system satisfied by *θ*_*n*_ allows us to provide a geometric characterization of the phase diagram for coupled gene circuits. The three distribution shapes correspond to three different regions in the phase diagram and the boundaries between them are characterized by two continuous increasing curves: a smooth curve *𝒞*_1_ and a piecewise-smooth curve *𝒞*_2_. Interestingly, we showed that the two curves can be constructed geometrically by the contour lines of *θ*_*n*_. The smooth curve *𝒞*_1_ is the contour line of *θ*_1_, and it separates the phase diagram into a region for bell-shaped distributions and the other region for decaying and bimodal distributions. The piecewise-smooth curve *𝒞*_2_ is the composed of the contour lines of *θ*_*n*_, *n ≥* 2, and it further separates the latter region into a subregion for decaying distributions and the other subregion for bimodal distributions. The bimodal parameter region is then enclosed by the two curves and the coordinate axes. Based on the geometric structure of the phase diagram, we provided some simple and verifiable sufficient and/or necessary conditions for the existence of the bimodal region, as well as some some simple and verifiable sufficient and/or necessary conditions for the steady-state distribution to be decaying, bell-shaped, or bimodal. In particular, we showed that bimodality can only occur when the parameter set falls into the “potential bimodal region” enclosed by two straight lines and the coordinate axes. This generalized a previous result that an unregulated gene can only exhibit bimodality when both the gene switching rates are smaller than the degradation rate [24].

Finally, we examine how the phase diagram is affected by the strengths of positive and feedback feedback. Overall, the bimodal parameter region enlarges (shrinks) as positive (negative) feedback becomes stronger. For positive feedback loops, the bimodal region always exists (under the mild condition of *s >* 2). However, for negative feedback loops, the bimodal region only exists when negative feedback is weak and may totally appear when negative feedback is very strong. For coupled feedback loops, the bimodal region may either enlarge or shrink as positive and negative feedback strengths increase simultaneously, depending on the ratio of the two feedback strengths.

Our model has two major limitations: (i) we assume that protein molecules are produced one at a time. However, many proteins are produced in a bursty manner due to rapid translation of protein from a single, short-lived mRNA molecule [42–44]. Hence our results are more applicable to the case where the mRNA lifetime is long (as common in mammalian cells [45]), where protein synthesis may appear non-bursty; (ii) we assume that there is no change in the protein number during gene activation and inactivation. However, in reality, the protein number decreases by one when a protein copy binds to a gene and increases by one when unbinding occurs [19, 20]. In other words, the present model ignores the protein-gene binding fluctuations and hence it may not be accurate when the protein number is very small or when the feedback strength is very strong [20, 46]. The reason why we make the above two approximations is that if we ignore protein bursting and binding fluctuations, then *θ*_*n*_ satisfies Eqs. (15) and (16) which are crucial to our geometric theory. However, if protein bursting and binding are taken into account, then these two equations are broken and an in-depth analysis will become much more difficult.

We anticipate to address the above two issues in our future work and to investigate the phase diagram in the presence of cooperative transcription regulation [13, 47] and non-exponential elongation or degradation [48, 49]. In addition, we also hope to develop the geometric theory of the dynamical phase diagram for time-dependent protein distributions [22, 50, 51].

## Acknowledgements

G. L. acknowledges support from National Natural Science Foundation of China with grant No. 12371162. F. J. acknowledges support from National Natural Science Foundation of China with grant No. 12271118. C. J. acknowledges support from National Natural Science Foundation of China with grant Nos. U2230402 and 12271020.

## Appendix A Proof of Theorem 4.4

Before proving Theorem 4.4, we give five lemmas whose proof can be found in Supplementary Sections 3-6.

### Lemma A.1.

Let *s >* 1, *µ, ν ≥* 0 be fixed and let *ω < s* be the constant given in Eq. (4). For any *n* ∈ [1, *s*), the equation *θ*_*n*_(*a, b*) = 1 determines a curve *a* = *g*_*n*_(*b*) in the *b*-*a* plane, which has the following properties:

i. *g*_*n*_(*b*) is defined for any 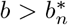, where 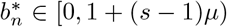. In particular, we have 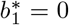 and 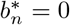 for any *n* ∈ [*ω, s*). For any 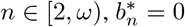 if and only if *θ*_*n*_(0, 0) = lim_(*a,b*)*→*(0,0)_ *θ*_*n*_(*a, b*) *≤* 1.
ii. *g*_1_(*b*) *<* (*s* + *b*)*/*(*s −* 1) and lim_*b→∞*_ *g*_*n*_(*b*)*/b* = *n/*(*s − n*) for any *n* ∈ [1, *s*).
iii. If *ν* = 0, then *g*_1_(0) = lim_*b→*0_+ *g*_1_(*b*) = 0 and 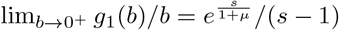.
iv. If *ν >* 0, then *g*_1_(0) = lim_*b→*0_+ *g*_1_(*b*) *>* 0 is finite. If *ω ≤* 1, then we have *ω ≤ g*_1_(0) *≤ s/*(*s −* 1); if *ω >* 1, then we have 0 *< g*_1_(0) *≤* min*{ω, s/*(*s −* 1)*}*. For any *n* ∈ [*ω, s*), we have *g*_*n*_(0) *≥ ω*; for any *n* ∈ [2, *ω*), we have

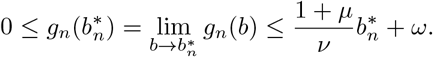

Recall the definition of *Q*_*n*_ given in Eq. (9). In the following, we rewrite *Q*_*n*_ as *Q*_*n*_(*a, b*) to emphasize its dependence on *a* and *b*. Note that *Q*_*n*_(*a, b*) = 0 defines a contour line of *Q*_*n*_, which is the straight line

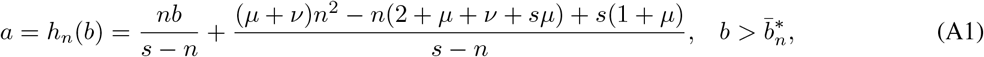

where

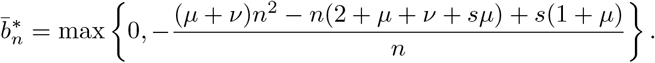

In particular, it is easy to see that 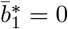 whenever *s >* 2.

### Lemma A.2.

Let *s >* 2 and *µ, ν ≥* 0 be fixed and let *n*_1_ be the constant given in Eq. (23). Let *a* = *h*_*n*_(*b*) be the contour line of *Q*_*n*_ defined above.

i. Then we have

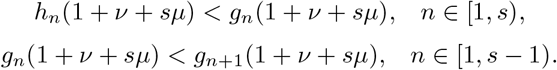
ii. For any *n* ∈ [*n*_1_, *s*), the two curves *a* = *g*_*n*_(*b*) and *a* = *h*_*n*_(*b*) do not intersect. In this case, we have 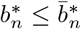 and *g*_*n*_(*b*) *> h*_*n*_(*b*). Moreover, for any *n* ∈ [*n*_1_, *s −* 1), the three curves *a* = *g*_*n*_(*b*), *a* = *h*_*n*_(*b*), and *a* = *g*_*n*+1_(*b*) do not intersect. In this case, we have 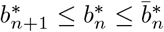 and *g*_*n*+1_(*b*) *> g*_*n*_(*b*) *> h*_*n*_(*b*).
iii. Let *n*_1_ *>* 1. If *a* = *g*_*n*_(*b*) and *a* = *h*_*n*_(*b*) intersect at *b* = *b*_*n*_ for some *n* ∈ [1, *n*_1_), then *a* = *g*_*n*_(*b*), *a* = *h*_*n*_(*b*), and *a* = *g*_*n*+1_(*b*) must intersect at the same point. In this case, we have 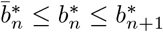 and

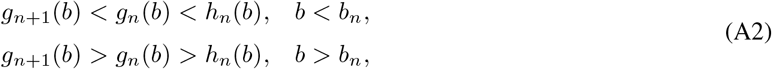

If *a* = *g*_*n*_(*b*) and *a* = *h*_*n*_(*b*) do not intersect for some *n* ∈ [1, *n*_1_), then *a* = *g*_*n*_(*b*), *a* = *h*_*n*_(*b*), and *a* = *g*_*n*+1_(*b*) do not intersect. In this case, we have 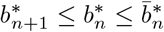 and *g*_*n*+1_(*b*) *> g*_*n*_(*b*) *> h*_*n*_(*b*).

### Lemma A.3.

Let *s >* 2 + 2*ν/*(1 + *µ*) and *µ, ν ≥* 0 be fixed. Then *n*_1_ *>* 1. Suppose that *a* = *g*_*n*_(*b*) and *a* = *h*_*n*_(*b*) intersect for some 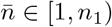. Then *a* = *g*_*n*_(*b*) and *a* = *h*_*n*_(*b*) also intersect for any 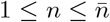. Let (*b*_*n*_, *a*_*n*_) be the intersection point of *a* = *g*_*n*_(*b*) and *a* = *h*_*n*_(*b*) in the *b*-*a* plane for each 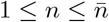. Then we have

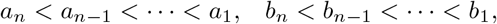

and

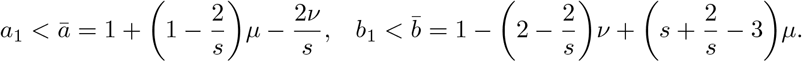

### Lemma A.4.

Let *s >* 2 and *µ, ν ≥* 0 be fixed. Let *n*_2_ be the largest integer in the interval [1, *s*).

i. If 2 *< s ≤* 2 + 2*ν/*(1 + *µ*), then we have 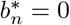 and *g*_1_(*b*) *< g*_*n*_(*b*), *b >* 0 for all *n* ∈ [2, *s*).
ii. If *s >* 2 + 2*ν/*(1 + *µ*) and 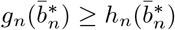 for some *n* ∈ [1, *s*), then we have 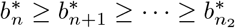 and

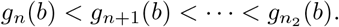

In particular, if *s >* 2 + 2*ν/*(1 + *µ*) and *g*_1_(0) *≥ h*_1_(0) = (*s −* 2)*/*(*s −* 1), then we have 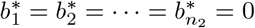 and

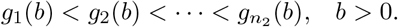

Given the above lemmas, we are now in a position to prove Theorem 4.4.

*Proof of Theorem 4*.*4*. When *s >* 1, Lemma A.1(i) shows that *θ*_1_(*a, b*) = 1 determines a curve *𝒞*_1_: *a* = *g*_1_(*b*), *b >* 0 in the *b*-*a* plane. The smoothness of *g*_1_(*b*) follows from the monotonicity of *θ*_1_(*a, b*), as proved in Lemma 4.3, and the implicit function theorem. Moreover, it is easy to see that

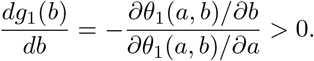

This shows that *g*_1_(*b*) increases strictly with respect to *b*. Hence *𝒞*_1_ is a smooth and strictly increasing curve (see the green curve in Fig. 4). By Lemma A.1(iii) and (iv), we have *g*_1_(0) *≥* 0 and *g*_1_(0) = 0 if and only if *ν* = 0. It follows from Lemma A.1(ii) and (iv) that *g*_1_(0) *≤ s/*(*s −* 1) and lim_*b→∞*_ *g*_1_(*b*)*/b* = 1*/*(*s −* 1). This proves Eq. (20). In addition, when *ν* = 0, it follows from Lemma A.1(iii) that

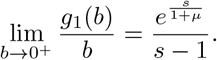

**Figure 4.**
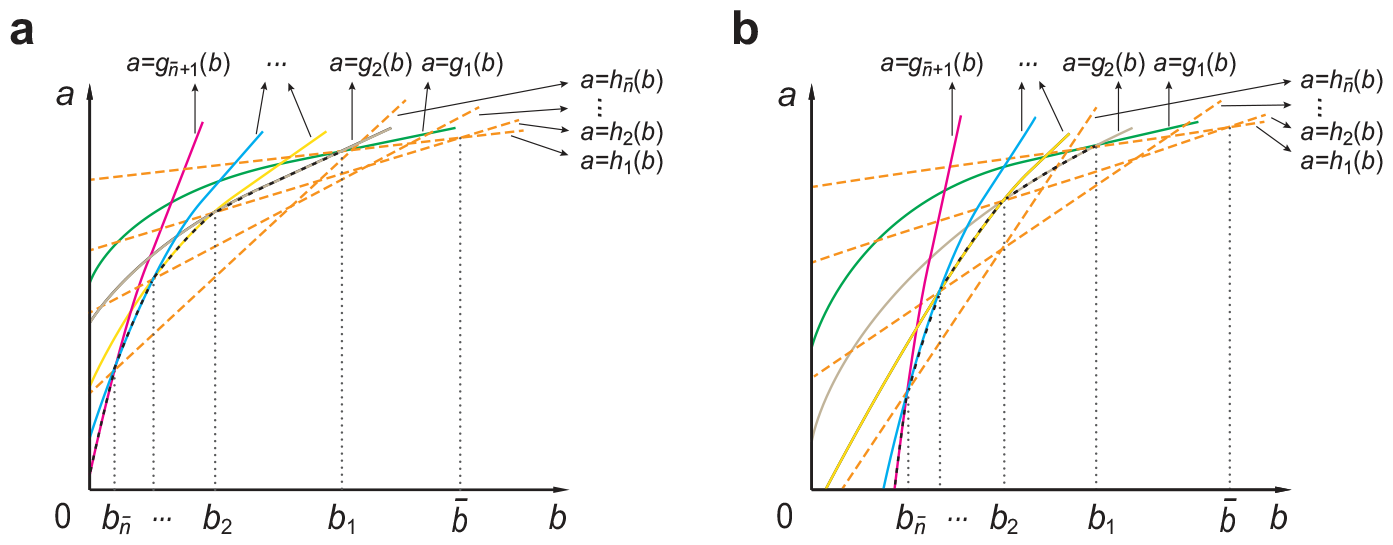
Construction of the curves *𝒞*_1_ and *𝒞*_2_ in the phase diagram. For each integer *n* ∈ [1, *s*), the equation *θ*_*n*_ = *P*_*n*_*/P*_*n−*1_ *≡* 1 determines a smooth and strictly increasing curve *a* = *g*_*n*_(*b*) (solid curves), and the equation *Q*_*n*_ = 0 determines the straight line *a* = *h*_*n*_(*b*) given by Eq. (A1) (yellow dashed lines). It can be proved that there exists 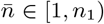 such that the three curves *a* = *h*_*n*_(*b*), *a* = *g*_*n*_(*b*), and *a* = *g*_*n*+1_(*b*) intersect at a unique point *b* = *b*_*n*_ for all 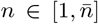, while they do not intersect for all 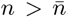. The straight lines *a* = *h*_1_(*b*) and *a* = *h*_2_(*b*) intersect at 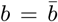, and the curve *𝒞*_1_ (green curve) is defined by *a* = *g*_1_(*b*) which intersects with *a* = *g*_2_(*b*) at 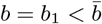. The curve *𝒞*_2_ (black dashed curve) is constructed by splicing the curves 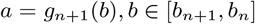 for 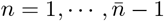 and the curve 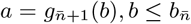. The curve *𝒞*_2_ may start from the *a*-axis as shown in **(a)** or start from the *b*-axis as shown in **(b)**.

When *ν >* 0, it follows from Lemma A.1(iv) that 0 *< g*_1_(0) *≤ s/*(*s −* 1). This shows that lim_*b→*0_+ *g*_1_(*b*)*/b* = *∞* and thus we obtain Eq. (21).

We now prove part (i) of Theorem 4.4 under the condition of *g*_1_(0) *≥* (*s −* 2)*/*(*s −* 1). First, Lemma A.1(ii) shows that *g*_1_(*b*) *<* (*s* + *b*)*/*(*s −* 1). This shows that *𝒞*_1_ lies below the straight line *L*_1_ : *a* = (*s* + *b*)*/*(*s −* 1), *b >* 0. To prove that *𝒞*_1_ lies above the straight line *L*_2_: *a* = (*s* + *b −* 2)*/*(*s −* 1), *b >* 0, note that the contour line *a* = *h*_1_(*b*) of *Q*_1_ defined in Eq. (A1) is exactly the straight line *L*_2_ (see the orange dashed line in Fig. 4). Substitute *n* = 1 and *a* = *g*_1_(*b*) into Eq. (12), we obtain

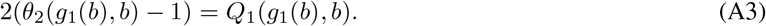

When 1 *< s ≤* 2, it follows from Lemma 4.1 that *θ*_2_(*a, b*) *< s/*2 *≤* 1 for all *a, b >* 0. This indicates that

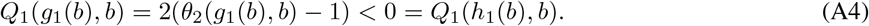

It is easy to see that *Q*_1_(*a, b*) strictly decreases with respect to *a* for any *s >* 1. This fact, together with Eq. (A4), shows that *g*_1_(*b*) *> h*_1_(*b*), *b >* 0. This shows that *𝒞*_1_ lies above *L*_2_ when 1 *< s ≤* 2.

We next prove that *𝒞*_1_ lies above *L*_2_ when *s >* 2. Lemma A.1 shows that *θ*_*n*_(*a, b*) = 1 determines a smooth and strictly increasing curve 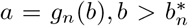 in the *b*-*a* plane for any *n* ∈ [2, *s*). When 2 *< s ≤* 2 + 2*ν/*(1 + *µ*), it follows from Lemma A.4(i) that 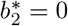 and *g*_1_(*b*) *< g*_2_(*b*), *b >* 0. When *s >* 2 + 2*ν/*(1 + *µ*), since we have assumed that *g*_1_(0) *≥* (*s −* 2)*/*(*s −* 1) = *h*_1_(0), it follows from Lemma A.4(ii) that 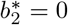 and *g*_1_(*b*) *< g*_2_(*b*), *b >* 0. Hence whenever *s >* 2, we have *g*_1_(*b*) *< g*_2_(*b*). Since *θ*_1_(*a, b*) is strictly increasing with respect to *a*, it follows from Eq. (A3) that

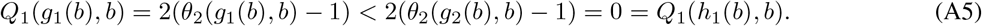

Since *Q*_1_(*a, b*) strictly decreases with respect to *a*, we have *g*_1_(*b*) *> h*_1_(*b*), *b >* 0. This shows that *𝒞*_1_ lies above *L*_2_ when *s >* 2.

We now prove that the parameter regions below and above *𝒞*_1_ correspond to decaying and bell-shaped distributions, respectively. Given a point (*b, a*) satisfying *a ≤ g*_1_(*b*). Since *θ*_1_(*a, b*) is strictly increasing with respect to *a*, we have

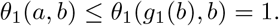

This shows that *P*_1_ *≤ P*_0_. When 1 *< s ≤* 2, we have *θ*_*n*_(*a, b*) *< s/n ≤* 1 for all *n* ∈ [2, *s*). This shows that *P*_*n*_ *< P*_*n−*1_ for all *n* ∈ [2, *s*). When *s >* 2, since we have assumed that *g*_1_(0) *≥* (*s −* 2)*/*(*s −* 1), it follows from Lemma A.4(i) and (ii) that *a ≤ g*_1_(*b*) *< g*_*n*_(*b*) for any *n* ∈ [2, *s*). Then the monotonicity of *θ*_*n*_(*a, b*) with respect to *a* yields

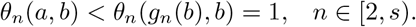

This shows that *P*_*n*_ *< P*_*n−*1_ for all *n* ∈ [2, *s*). Moreover, it follows from Lemma 4.1 that we have *θ*_*n*_(*a, b*) *< s/n ≤* 1 when *n ≥ s*. This shows that *P*_*n*_ *< P*_*n−*1_ for any *n ≥ s*. Hence *P*_*n*_ is decaying whenever *a ≤ g*_1_(*b*), i.e. in the parameter region below or on *𝒞*_1_. On the other hand, given a point (*b, a*) satisfying *a > g*_1_(*b*). Since *θ*_*n*_(*a, b*) is strictly increasing with respect to *a*, we have

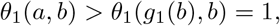

and hence *P*_1_ *> P*_0_. Note that among decaying, bell-shaped, and bimodal distributions, only bell-shaped distributions give rise to *P*_1_ *> P*_0_. Hence *P*_*n*_ is bell-shaped whenever *a > g*_1_(*b*), i.e. in the parameter region above *𝒞*_1_.

Finally we prove part (ii) of Theorem 4.4 under the condition of *g*_1_(0) *<* (*s −* 2)*/*(*s −* 1). Since *g*_1_(0) *≥* 0, we have *s >* 2. When 2 *< s ≤* 2 + 2*ν/*(1 + *µ*), it follows from Lemma A.4(i) that 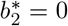 and *g*_1_(*b*) *< g*_2_(*b*), *b >* 0. It thus follows from Eq. (A5) that *g*_1_(*b*) *> h*_1_(*b*), *b >* 0. This indicates that

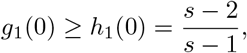

which contradicts our assumption of *g*_1_(0) *<* (*s −* 2)*/*(*s −* 1). Hence we must have *s >* 2 + 2*ν/*(1 + *µ*), i.e. *ω >* 2.

By Lemma A.2(i), we have *g*_1_(1 + *ν* + *sµ*) *> h*_1_(1 + *ν* + *sµ*). Since we have assumed that *g*_1_(0) *≥ h*_1_(0), the curves *a* = *g*_1_(*b*) and *a* = *h*_1_(*b*) must intersect at some point (*b, a*) = (*b*_1_, *a*_1_). Since *ω >* 2, we have *n*_1_ *≥ ω −* 1 *>* 1 and thus Lemma A.2(iii) shows that *a* = *g*_1_(*b*) and *a* = *h*_1_(*b*) must intersect at a unique point (otherwise Eq. (A2) will be broken). In other words, *𝒞*_1_ intersects with *L*_2_ at a unique point. By Lemma A.3, the intersection point (*b*_1_, *a*_1_) satisfies Eq. (22). Furthermore, by Lemma A.2(iii), we have

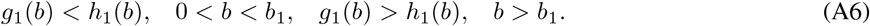

Hence *𝒞*_1_ lies below *L*_2_ for *b < b*_1_ and lies above *L*_2_ for *b > b*_1_. Recall that we have proved that *𝒞*_1_ lies below *L*_1_; this shows that *𝒞*_1_ lies between *L*_1_ and *L*_2_ for *b > b*_1_.

We now construct the curve *𝒞*_2_. Let 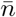 be the largest integer in the interval [1, *n*_1_) such that the curves 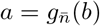 and 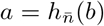 intersect. Since 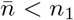, it is easy to verify that 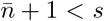. By Lemma A.2(iii) and Lemma A.3, the curves *a* = *g*_*n*_(*b*), *a* = *h*_*n*_(*b*), and *a* = *g*_*n*+1_(*b*) must intersect at a unique point (*b*_*n*_, *a*_*n*_) for all 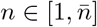. It follows from Lemma A.2(iii) that

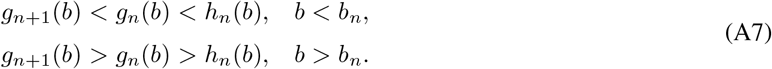

and Lemma A.3 further shows that

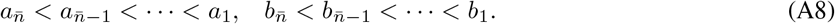

If 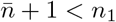, then by the choice of 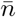, it is clear that 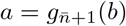 and 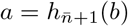 do not intersect. If 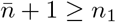, then it follows from Lemma A.2(ii) that 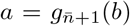 and 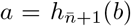 also do not intersect. In both situations, it follows from Lemma A.2(ii) and (iii) that 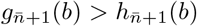. Hence it follows from Lemma A.4(ii) that

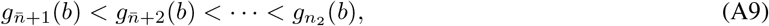

where *n*_2_ is the largest integer in the interval [1, *s*). Hence 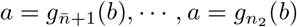 do not intersect. Hence the curve *𝒞*_2_ can be constructed by the the piecewise smooth, strictly increasing curve *a* = *g*(*b*) (see the black dashed curve in Fig. 4) with

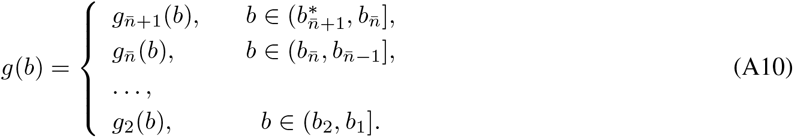

From Eqs. (A7), (A8), and (A9), it is easy to see that *a* = *g*(*b*) intersects with *a* = *g*_1_(*b*) at *b* = *b*_1_ and *g*(*b*) *≤ g*_*n*_(*b*) for all *n* ∈ [1, *s*). Here it may occur that 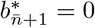 (see Fig. 4(a) for an illustration) or 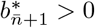 (see Fig. 4(b) for an illustration). Since *g*(*b*) *< g*_1_(*b*), it is clear that *𝒞*_2_ lies below *𝒞*_1_ for *b < b*_1_. In addition, since *a* = *g*_1_(*b*), *a* = *h*_1_(*b*), and *a* = *g*_2_(*b*) intersect at *b* = *b*_1_, it is clear that *𝒞*_2_ intersects with *𝒞*_1_ and *L*_2_ at a unique point (*b, a*) = (*b*_1_, *a*_1_).

We next analyze the distribution shapes in the three different parameter regions separated by *𝒞*_1_ and *𝒞*_2_. Given a point (*b, a*) satisfying *a ≥ g*_1_(*b*) with 0 *< b < b*_1_, Eq. (A7) shows that *a ≥ g*_1_(*b*) *> g*_2_(*b*). Since *θ*_*n*_(*a, b*) is strictly increasing with respect to *a*, we have

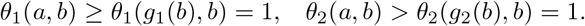

This indicates that *P*_0_ *≤ P*_1_ *< P*_2_. Hence in this case, *P*_*n*_ is bell-shaped. On the other hand, given a point (*b, a*) satisfying *a > g*_1_(*b*) with *b ≥ b*_1_, by the monotonicity of *θ*_*n*_(*a, b*) with respect to *a*, we have

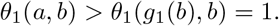

This shows that *P*_1_ *> P*_0_. In this case, *P*_*n*_ also is bell-shaped. Taken together, we have proved that *P*_*n*_ is bell-shaped whenever *a > g*_1_(*b*), i.e. in the parameter region above or on *𝒞*_1_.

Similarly, given a point (*b, a*) satisfying *a ≤ g*(*b*) with 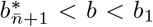 (the parameter region below *𝒞*_2_ and on the left-hand side of *b* = *b*_1_). We have proved that we have *g*(*b*) *≤ g*_*n*_(*b*) for all *n* ∈ [1, *s*). Since *θ*_*n*_(*a, b*) is strictly increasing with respect to *a*, we have

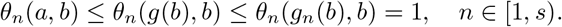

This shows that *P*_*n*_ *≤ P*_*n−*1_ for all *n* ∈ [1, *s*). Moreover, Lemma 4.1 indicates that *θ*_*n*_ *< s/n ≤* 1 for all *n ≥ s*. Hence in this case, *P*_*n*_ is decaying. On the other hand, given a point (*b, a*) satisfying *a ≤ g*_1_(*b*) with *b ≥ b*_1_ (the parameter region below *𝒞*_1_ and on the right-hand side of *b* = *b*_1_), it follows from Lemma A.2(iii) and Lemma A.3 that

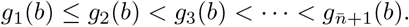

This fact, together with Eq. (A9), shows that *g*_1_(*b*) *≤ g*_*n*_(*b*) for all *n* ∈ [2, *s*). Since *θ*_*n*_(*a, b*) is strictly increasing with respect to *a*, we have

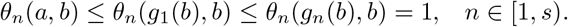

This shows that *P*_*n*_ *≤ P*_*n−*1_ for all *n* ∈ [1, *s*). Moreover, Lemma 4.1 indicates that *θ*_*n*_ *< s/n ≤* 1 for all *n ≥ s*. Hence in this case, *P*_*n*_ is also decaying. Taken together, we have proved that *P*_*n*_ is decaying in the parameter region below *𝒞*_1_ and on the right-hand side of *𝒞*_2_.

We finally consider the rest of the parameter region surrounded by *𝒞*_1_, *𝒞*_2_, and the coordinate axes. If *𝒞*_2_ starts\ from the *a*-axis, i.e. 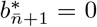, then we consider a point (*b, a*) satisfying *g*(*b*) *< a < g*_1_(*b*) for 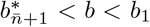. The monotonicity of *θ*_*n*_(*a, b*) with respect to *a* yields

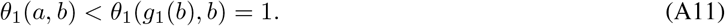

This shows that *P < P*. The definition of *g*(*b*) implies that there exists 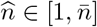 such that 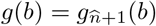. Then we have

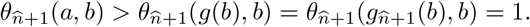

This shows that 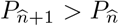. Hence in this case, *P*_*n*_ is bimodal. If *𝒞*_2_ starts from the *b*-axis, i.e. 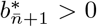, then we consider a point (*b, a*) satisfying *g*(*b*) *< a < g*_1_(*b*) for 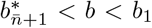 (the parameter region surrounded by *𝒞*_1_, *𝒞*_2_, and 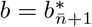), or satisfying *a < g*_1_(*b*) for 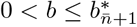 (the parameter region surrounded by 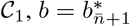, and the coordinate axes). For the former region, similar arguments show that *P*_*n*_ is bimodal. For the latter region, it follows from Eq. (A11) that *P*_1_ *< P*_0_. By the definition of 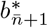, we have 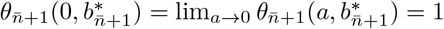. Then the monotonicity of *θ*_*n*_(*a, b*) with respect to *a* and *b* yields

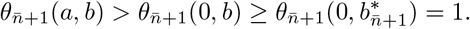

This shows that 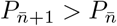. Hence in this case, *P*_*n*_ is also bimodal. Taken together, we have proved that *P*_*n*_ is bimodal in the parameter region surrounded by *𝒞*_1_, *𝒞*_2_, and the coordinate axes. The proof is completed.

