## supplemental materials for "Geometry theory of distribution shapes for autoregulatory gene circuits"

#### Contents

**1 Proof of Lemma 4.1** **2**

**2 Proof of Eq. (25) in the main text** **3**

**3 Proof of Lemma A.1** **4**

**4 Proof of Lemma A.2** **9**

**5 Proof of Lemma A.3** **16**

**6 Proof of Lemma A.4** **17**

### 1 Proof of Lemma 4.1

**Lemma 4.1.** For any $s, a, b > 0$ and $\mu, \nu \geq 0$, we have

$$\theta_n = \frac{P_n}{P_{n-1}} < \frac{s}{n}, \quad n \geq 1.$$

*Proof.* Firstly, we prove $\lim_{k\to\infty} \theta_n(a + k(1+\mu), b + k\nu) = \omega/n$, where $\omega = s(1+\mu)/(1+\mu+\nu)$. Noticing $\alpha = a/(1+\mu)$ and $\beta = (a+b)/(1+\mu+\nu) + s\nu(1+\mu+\nu)^2$, we find

$$\frac{(\alpha+n+m)\cdots(\alpha+n+m+k-1)(1+\mu)^k}{(\beta+n+m)\cdots(\beta+n+m+k-1)(1+\mu+\nu)^k} < 1.$$

Thus

$$\begin{aligned}
&|{}_1F_1(\alpha+m+n; \beta+m+n; -\omega z_0)| \\
&= \left|1 + \sum_{k=1}^{\infty} \left[\frac{(\alpha+m+n)\cdots(\alpha+m+n+k-1)}{(\beta+m+n)\cdots(\beta+m+n+k-1)} \cdot \frac{(-\omega z_0)^k}{k!}\right]\right| \\
&< 1 + \sum_{k=1}^{\infty} \left[\frac{(\alpha+m+n)\cdots(\alpha+m+n+k-1)(1+\mu)^k}{(\beta+m+n)\cdots(\beta+m+n+k-1)(1+\mu+\nu)^k} \cdot \frac{(sz_0)^k}{k!}\right] \\
&< 1 + \sum_{k=1}^{\infty} \frac{(sz_0)^k}{k!} = e^{sz_0},
\end{aligned} \tag{1}$$

where $z_0 = 1/(1+\mu+\nu)$. By Lebesgue dominated convergence theorem, we find that the limit of ${}_1F_1(\alpha+m+n; \beta+m+n; -\omega z_0)$ exsits when $m \to \infty$ for $n \geq 1$. Therefore, the limit of

$$\theta_n(a + m(1+\mu), b + m\nu) = \frac{\alpha+m+n-1}{\beta+m+n-1} \cdot \frac{\omega}{n} \cdot \frac{{}_1F_1(\alpha+m+n; \beta+m+n; -\omega z_0)}{{}_1F_1(\alpha+m+n-1; \beta+m+n-1; -\omega z_0)}$$

exists when $m \to \infty$ for $n \geq 1$, and

$$\begin{aligned}
&\lim_{m\to\infty} \theta_n(a + m(1+\mu), b + m\nu) \\
&= \lim_{m\to\infty} \left[\frac{\alpha+m+n-1}{\beta+m+n-1} \cdot \frac{\omega}{n} \cdot \frac{{}_1F_1(\alpha+m+n; \beta+m+n; -\omega z_0)}{{}_1F_1(\alpha+m+n-1; \beta+m+n-1; -\omega z_0)}\right] \\
&= \lim_{m\to\infty} \left[\frac{\alpha+m+n-1}{\beta+m+n-1} \cdot \frac{\omega}{n} \cdot \frac{1 + \sum_{k=1}^{\infty} \left[\frac{(\alpha+n+m)\cdots(\alpha+m+n+k-1)}{(\beta+m+n)\cdots(\beta+m+n+k-1)} \frac{(-\omega z_0)^k}{k!}\right]}{1 + \sum_{k=1}^{\infty} \left[\frac{(\alpha+m+n-1)\cdots(\alpha+m+n+k-2)}{(\beta+m+n-1)\cdots(\beta+m+n+k-2)} \frac{(-\omega z_0)^k}{k!}\right]}\right] = \frac{\omega}{n}.
\end{aligned} \tag{2}$$

Then we try to prove $\theta_n = P_n/P_{n-1} < s/n$. When $\nu = 0$, we find $\alpha = a/(1+\mu) < \beta = (a+b)/(1+\mu)$ for all $a, b, s > 0$ and $\mu \geq 0$. Then the following equation holds:

$$P_n = \frac{s^n}{n!} \frac{\int_0^1 e^{-sz_0 t} t^{\alpha+n-1}(1-t)^{\beta-\alpha-1} dt}{\int_0^1 e^{s(1-z_0)t} t^{\alpha-1}(1-t)^{\beta-\alpha-1} dt}.$$

Thus for all $n \geq 1$, we have

$$\frac{P_n}{P_{n-1}} = \frac{s}{n} \frac{\int_0^1 e^{-sz_0 t} t^{\alpha+n-1}(1-t)^{\beta-\alpha-1} dt}{\int_0^1 e^{-sz_0 t} t^{\alpha+n-2}(1-t)^{\beta-\alpha-1} dt} < \frac{s}{n}.$$

Next we consider $\nu > 0$. By

$$\theta_n(a+1+\mu, b+\nu) = \frac{\alpha+n}{\beta+n} \cdot \frac{\omega}{n} \cdot \frac{{}_1F_1(\alpha+n+1; \beta+n+1; -\omega z_0)}{{}_1F_1(\alpha+n; \beta+n; -\omega z_0)} = \frac{n+1}{n} \theta_{n+1}(a, b), \tag{3}$$

we can obtain

$$\theta_{n+1}(a,b) = \frac{n}{n+1}\theta_n(a+1+\mu, b+\nu).$$

Replacing $n$ by $n+m-1$ in the above equation gives

$$\begin{aligned}\theta_{n+m}(a,b) &= \frac{n+m-1}{n+m}\theta_{n+m-1}(a+1+\mu, b+\nu) \\ &= \cdots = \frac{n}{n+m}\theta_n(a+m(1+\mu), b+m\nu), \quad m \geq 0.\end{aligned} \tag{4}$$

This indicates

$$\frac{n+m}{n}\theta_{n+m}(a,b) = \theta_n(a+m(1+\mu), b+m\nu), \quad m \geq 0.$$

Combining the above equation with Eq. (2), we have

$$\lim_{m\to\infty}\left[\frac{n+m}{n}\theta_{n+m}(a,b)\right] = \lim_{m\to\infty}\theta_n(a+m(1+\mu), b+m\nu) = \frac{\omega}{n}, \quad n \geq 1. \tag{5}$$

Then we suppose for contradiction that there exists $n' \geq 1$ such that $P_{n'}/P_{n'-1} \geq s/n'$. We verify the following equation in the text:

$$n(n+1)(\theta_{n+1}-1) = s[(n-1)(1+\mu)+a](1-1/\theta_n) + Q(n), \tag{6}$$

where

$$Q(n) = n^2(\mu+\nu) + n(a-1-\mu+b-1-\nu-s\mu) - s(a-1-\mu). \tag{7}$$

Taking $P_{n'}/P_{n'-1} \geq s/n'$ into Eq. (6), we obtain

$$\theta_{n'+1}(a,b) \geq \frac{(n'-1)\nu+b+s}{n'+1} > \frac{s}{n'+1}.$$

Repeating this process, we obtain $P_n/P_{n-1} > s/n$ for all $n > n'$. Thus

$$\frac{n'+m}{n'}\theta_{n'+m}(a,b) > \frac{s}{n'}$$

with $m \geq 1$ and

$$\lim_{m\to\infty}\left[\frac{n'+m}{n'}\theta_{n'+m}(a,b)\right] \geq \frac{s}{n'} > \frac{\omega}{n'}.$$

It contradicts Eq. (5), so $P_n/P_{n-1} < s/n$ for all $n \geq 1$. The proof is completed. □

### 2 Proof of Eq. (25) in the main text

When $a, b, \nu \ll 1$, the Markov chain shown in Fig. 1(b) in the main text has two different time scales and hence can be simplified using standard two-time-scale simplification techniques for Markov chains [1]. Since $\mu$ and $d = 1$ are large (relatively to $a, b, \nu$), all the microstates $(0, n)$ with $n \geq 1$ (often called the “fast states") will rapidly transition to the other microstates (often called the “slow states"). Once the system has entered the slow states, it will rarely transition back to the fast states because the transition rates from the slow states to the fast states are very small. According to the decimation theory for Markov chains [2, 3], all the fast states can be removed from the system and thus the original chain can be simplified to the reduced chain illustrated in Supplementary Fig. 1. The remaining question is how to

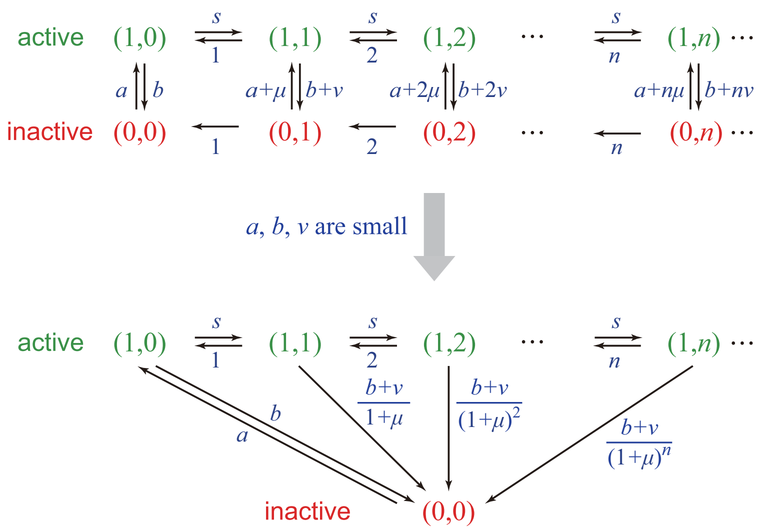


Figure 1: **Model reduction.** When $a, b, \nu \ll 1$, using the decimation theory [2, 3], the Markov chain illustrated in the upper panel can be reduced to be one illustrated in the lower panel.

determine the effective transition rates of the reduced chain. According to the decimation theory, once microstates $(0, n)$ with $n \geq 1$ are removed, there is an effective transition rate from microstate $(1, n)$ with $n \geq 1$ to microstate $(0, 0)$, i.e.

$$\tilde{q}_{(1,n),(0,0)} = q_{(1,n),(0,n)} p_{(0,n),(0,n-1)} \cdots p_{(0,1),(0,0)}, \quad n \geq 1.$$

Here $\tilde{q}_{x,y}$ represents the effective transition rate from microstate $x$ to microstate $y$ for the reduced chain, $q_{x,y}$ represents the transition rate from $x$ to $y$ for the original chain, and

$$p_{x,y} = \frac{q_{x,y}}{\sum_{z \neq y} q_{x,z}}$$

represents the probability weight from $x$ to $y$ (see [2, 3] for a detailed explanation). Note that $q_{(1,n),(0,n)} = b + n\nu$ and

$$p_{(0,n),(0,n-1)} = \frac{nd}{nd + b + n\mu} \approx \frac{1}{1 + \mu}.$$

Hence the effective transition rate from microstate $(1, n)$ with $n \geq 1$ to microstate $(0, 0)$ is given by

$$\tilde{q}_{(1,n),(0,0)} = (b + n\nu) \left( \frac{1}{1 + \mu} \right)^n, \quad n \geq 1.$$

Thus far, we have obtained all effective transition rates for the reduced chain. It is easy to prove that the steady-state protein distribution for the reduced chain is the ZIP distribution given in Eq. (25) in the main text.

### 3 Proof of Lemma A.1

To prove Lemma A.1, we need to prove the following lemma.

**Lemma 1.** (i) If $a \leq 1 + \mu$ and $a \geq -b + 2 + \mu + \nu + s\mu$, then $P_{n-1} > P_n$ for all $n \geq 1$;

(ii) If $s > 1$, $a \geq 1 + \mu$ and $a \leq \frac{b}{s-1} + \frac{s-2}{s-1}$, then $P_n$ is a decaying distribution.

*Proof.* (i) Taking $a-1-\mu+b-1-\nu-s\mu \geq 0$ and $a \leq 1+\mu$ into Eq. (7), we have $Q_n > 0$ for all $n \geq 1$ or $Q_n = 0$ for all $n \geq 1$. First, we consider $Q_n = 0$ for all $n \geq 1$. Noticing $Q_n = 0$ for all $n \geq 1$ if and only if $\mu = \nu = 0$, $a-1-\mu+b-1-\nu-s\mu = 0$ and $a = 1+\mu$. Taking $\mu = \nu = 0$, $a-1-\mu+b-1-\nu-s\mu = 0$ and $a = 1+\mu$ into

$$n(n+1)\Delta P_{n+1} = s[(n-1)(1+\mu)+a]\Delta P_n + Q(n)P_n, \quad n \geq 1 \tag{8}$$

which is proved in the text, we find

$$n(n+1)\Delta P_{n+1} = \frac{s}{n+1}\Delta P_n.$$

If $\Delta P_n \geq 0$, then $\Delta P_{n+1} \geq 0$. This implies $P_n \geq P_{n-1}$ for all $n \geq 1$. However we have $\theta_n = P_n/P_{n-1} < s/n$ by Lemma 4.1, so $P_n < P_{n-1}$ for $n > s$. This gives a contradiction, so $P_n < P_{n-1}$ when $Q_n = 0$ for all $n \geq 1$. Next we consider $Q_n > 0$ for all $n \geq 1$. We give the proof by contradiction. Noticing that $P_n$ peaks twice at most, with possible modes at $n = 0$ and $n = n_1 > 0$, so we can suppose there exists $n^* \geq 1$ such that

$$\Delta P_{n^*} = P_{n^*} - P_{n^*-1} \geq 0, \quad \Delta P_{n^*+1} = P_{n^*+1} - P_{n^*} \leq 0,$$

which implies $Q(n^*) \leq 0$ by Eq. (8). This gives a contradiction. The proof is completed.

(ii) When $s > 1$, we find $Q(1) \geq 0$ indicates $a \leq \frac{b}{s-1} + \frac{s-2}{s-1}$, and $Q(0) = s(1+\mu-a) \leq 0$ gives $a \geq 1+\mu$. We suppose for contradiction that there exists a peak at $n = n^* \geq 1$ such that

$$\Delta P_{n^*} = P_{n^*} - P_{n^*-1} > 0, \quad \Delta P_{n^*+1} = P_{n^*+1} - P_{n^*} \leq 0,$$

which implies $Q(n^*) < 0$ by Eq. (8). However, when $Q(1) \geq 0$ and $Q(0) \leq 0$, we have $Q(n) \geq 0$ for all $n \geq 1$, which gives a contradiction. The proof is completed. □

Next we prove Lemma A.1.

**Lemma A.1.** Let $s > 1$, $\mu, \nu \geq 0$ be fixed. For any $n \in [1, s)$, the equation $\theta_n(a, b) = 1$ determines a curve $a = g_n(b)$ in the $b$-$a$ plane, which has the following properties:

(i) $g_n(b)$ is defined for any $b > b_n^*$, where $b_n^* \in [0, 1+(s-1)\mu)$. In particular, we have $b_1^* = 0$ and $b_n^* = 0$ for any $n \in [\omega, s)$. For any $n \in [2, \omega)$, $b_n^* = 0$ if and only if $\theta_n(0,0) = \lim_{(a,b)\to(0,0)} \theta_n(a, b) \leq 1$. Here $\omega = s(1+\mu)/(1+\mu+\nu)$.

(ii) $g_1(b) < (s+b)/(s-1)$ and $\lim_{b\to\infty} g_n(b)/b = n/(s-n)$ for any $n \in [1, s)$.

(iii) If $\nu = 0$, then $g_1(0) = \lim_{b\to 0^+} g_1(b) = 0$ and $\lim_{b\to 0^+} g_1(b)/b = e^{\frac{s}{1+\mu}}/(s-1)$.

(iv) If $\nu > 0$, then $g_1(0) = \lim_{b\to 0^+} g_1(b) > 0$ is finite. If $\omega \leq 1$, then we have $\omega \leq g_1(0) \leq s/(s-1)$; if $\omega > 1$, then we have $0 < g_1(0) \leq \min\{\omega, s/(s-1)\}$. For any $n \in [\omega, s)$, we have $g_n(0) \geq \omega$; for any $n \in [2, \omega)$, we have

$$0 \leq g_n(b_n^*) = \lim_{b \to b_n^*} g_n(b) \leq \frac{1+\mu}{\nu} b_n^* + \omega.$$

*Proof.* (i) Noticing that Lemma 4.1 indicates $\theta_n = P_n/P_{n-1} < s/n \leq 1$ for all $n \geq s$, so we only consider $n < s$. For each given $b > 0$, we know that when $a \to \infty$, $P_n$ tends to the Poisson distribution $P_n^* = s^n e^{-s}/n!$. This implies $\theta_n(a, b) \to s/n > 1$ as $a \to \infty$ for $1 \leq n < s$, i.e., there is sufficiently large $a$ such that $\theta_n(a, b) > 1$ for any given $b > 0$ and $n \geq 1$. If we can prove that there is some $a$ such that $\theta_n(a, b) < 1$ for any given $b$, then there is exactly one a to make $\theta_n(a, b) = 1$ by the monotonicity of $\theta_n(a, b)$ in $a$ and $b$ which is proved by Lemma 4.3 in the text, and so $a = g_n(b)$ is well defined for all $1 \leq n < s$. Thus we only need to prove that there exists some $a$ such that $\theta_n(a, b) < 1$ for any given $b$.

For $n = 1$, combining Eq. (1) with Lebesgue dominated convergence theorem, we find the limit of $\theta_1(a, b)$ exists for any given $b > 0$ when $a \to 0$ and

$$\lim_{a\to 0} \theta_1(a,b) = \lim_{a\to 0}\left[\frac{\alpha}{\beta}\cdot s\cdot \frac{1+\sum_{k=1}^{\infty}\frac{(\alpha+1)\dots(\alpha+k)}{(\beta+1)\dots(\beta+k)}\frac{(\frac{-s}{1+\mu})^k}{k!}}{1+\sum_{k=1}^{\infty}\frac{(\alpha)\dots(\alpha+k-1)}{(\beta)\dots(\beta+k-1)}\frac{(\frac{-s}{1+\mu})^k}{k!}}\right] = 0.$$

Define $\theta_1(0, b) = \lim_{a\to 0} \theta_1(a, b)$, then we have $\theta_1(0, b) = 0 < 1$ for any given $b > 0$. This indicates that there exists sufficiently small $a$ such that $\theta_1(a, b) < 1$ for any given $b > 0$. Therefore, $a = g_1(b)$ is well defined for all $b > 0$.

For $2 \le n < s$, when $a = 1 + \mu$ and $b = 1 + (s-1)\mu$, the conditions of Lemma 1 (ii) are met, so $P_n$ is decaying distribution, i.e.,

$$\theta_n(1 + \mu, 1 + (s-1)\mu) \le 1.$$

According to the monotonicity of $\theta_n(a, b)$ with respect to $a$ and $b$, we have

$$\theta_n(0, b) < \theta_n(1 + \mu, b) \le \theta_n(1 + \mu, 1 + (s-1)\mu) \le 1, \quad b \ge 1 + (s-1)\mu. \tag{9}$$

Combining Eq. (1) with Lebesgue dominated convergence theorem again, we find for $n \ge 1$

$$\begin{aligned}\lim_{a\to 0, b\to 0} \theta_n(a,b) &= \lim_{a\to 0, b\to 0}\left[\frac{\alpha+n-1}{\beta+n-1}\cdot\frac{\omega}{n}\cdot \frac{1+\sum_{k=1}^{\infty}\frac{(\alpha+n)\dots(\alpha+n+k-1)}{(\beta+n)\dots(\beta+n+k-1)}\frac{(-\omega z_0)^k}{k!}}{1+\sum_{k=1}^{\infty}\frac{(\alpha+n-1)\dots(\alpha+n+k-2)}{(\beta+n-1)\dots(\beta+n+k-2)}\frac{(-\omega z_0)^k}{k!}}\right]\\ &= \frac{n-1}{s\nu z_0 + n - 1}\cdot\frac{\omega}{n}\cdot\frac{F(n; s\nu z_0^2 + n; -\omega z_0)}{F(n-1; s\nu z_0^2 + n - 1; -\omega z_0)}\end{aligned} \tag{10}$$

Define $\theta_n(0, 0) = \lim_{a\to 0, b\to 0} \theta_n(a, b)$. We discuss the cases of $\nu = 0$ and $\nu > 0$, respectively. Taking $\nu = 0$ into Eq. (10), we obtain $\theta_n(0, 0) = s/n > 1$, where $2 \le n < s$. Combining this with Eq. (9), then the monotonicity of $\theta_n(a, b)$ with respect to $b$ imply that there exists $0 < b_n^* < 1 + (s-1)\mu$ such that $\theta_n(0, b_n^*) = 1$, and so

$$\theta_n(0, b) < 1, \quad b > b_n^*,$$

where $2 \le n < s$. This implies that there exists sufficiently small $a$ such that $\theta_n(a, b) < 1$ for any given $b > b_n^*$, so $a = g_n(b)$ is well defined for $b > b_n^*$ and $2 \le n < s$ when $\nu = 0$, where $b_n^* \in (0, 1 + (s-1)\mu)$.

When $\nu > 0$, it is obvious that $n < s\nu z_0^2 + n$ and $n - 1 < s\nu z_0^2 + n - 1$, so Eq. (10) can be rewritten as

$$\theta_n(0,0) = \frac{\omega}{n}\frac{\int_0^1 e^{-\omega z_0 t} t^{n-1}(1-t)^{s\nu z_0^2 - 1}\mathrm{d}t}{\int_0^1 e^{-\omega z_0 t} t^{n-2}(1-t)^{s\nu z_0^2 - 1}\mathrm{d}t} < \frac{\omega}{n}, \quad 2 \le n < s.$$

We find $\theta_n(0, 0) < 1$ when $\omega \le n < s$. Then we consider the cases of $\omega \le n < s$ and $2 \le n < \omega$, respectively. When $\omega \le n < s$, since $\theta_n(a, b)$ strictly decreases with respect to $b$, we have

$$\theta_n(0, b) < \theta_n(0, 0) < 1$$

for any given $b > 0$. This indicates that there is sufficiently small $a$ such that $\theta_n(a, b) < 1$, $\omega \le n < s$, for any given $b > 0$. Therefore, we obtain $a = g_n(b)$ is well defined for all $b > 0$ when $\omega \le n < s$ and $\nu > 0$.

Finally, we consider $2 \le n < \omega$ and $\nu > 0$. Since we can not know the size relationship between $\theta_n(0, 0)$ and 1 when $2 \le n < \omega$, we discuss the cases of $\theta_n(0, 0) \ge 1$ and $\theta_n(0, 0) < 1$, respectively, when $2 \le n < \omega$. If $\theta_n(0, 0) \le 1$, then we have

$$\theta_n(0, b) < \theta_n(0, 0) \le 1$$

by the monotonicity of $\theta_n(a,b)$ in $b$, which implies that there exists sufficiently small $a$ such that $\theta_n(a,b) < 1$ for any given $b > 0$. If $\theta_n(0,0) > 1$, combining with Eq. (9), then there exists $0 < b_n^* < 1 + (s-1)\mu$ such that $\theta_n(0, b_n^*) = 1$, and so

$$\theta_n(0,b) < 1, \quad b > b_n^*.$$

This implies that there exists sufficiently small $a$ such that $\theta_n(a,b) < 1$ for any given $b > b_n^*$, so $a = g_n(b)$ is well defined for $b > b_n^*$. Therefore, $a = g_n(b)$ is well defined for $b > b_n^*$, where $b_n^* = 0$ if $\theta_n(0,0) \le 1$ and $b_n^* \in (0, 1+(s-1)\mu)$ if $\theta_n(0,0) > 1$, when $2 \le n < \omega$ and $\nu > 0$. Then we conclude that $a = g_n(b)$ is well defined for $b > b_n^*$, where $b_n^* = 0$ if $\theta_n(0,0) \le 1$ and $b_n^* \in (0, 1+(s-1)\mu)$ if $\theta_n(0,0) > 1$ when $2 \le n < s$. Since $\theta_n(0,0) < 1$ when $\omega \le n < s$, we obtain $b_n^* = 0$ when $\omega \le n < s$.

The differentiability of $g_n(b)$ follows from the monotonicity of $\theta_n(a,b)$ in $a$ and $b$ proved by Lemma 4.3 in the text and the implicit function theorem for all $1 \le n < s$. It also satisfies

$$\frac{dg_n(b)}{db} = -\frac{\partial\theta_n(a,b)/\partial b}{\partial\theta_n(a,b)/\partial a} > 0, \quad 1 \le n < s,$$

thus $g_n(b)$ increases strictly with respect to $b$ for all $1 \le n < s$.

(ii) By the recurrence relation between $P_{n-1}, P_n$, and $P_{n+1}$

$$n(n+1)P_{n+1} - n[(n-1)(1+\mu+\nu) + a + b + s]P_n + s[(n-1)(1+\mu) + a]P_{n-1} = 0, \quad n \ge 1. \tag{11}$$

which is proved by Lemma 3.1 in the text, we obtain

$$(a+b+s)P_1 - saP_0 > 0$$

and so

$$\theta_1 = \frac{P_1}{P_0} > \frac{sa}{a+b+s}.$$

Then taking $a = g_1(b)$ into the above relation, we obtain

$$1 = \theta_1(g_1(b), b) > \frac{sg_1(b)}{g_1(b) + b + s}.$$

Therefore, we have $g_1(b) < b/(s-1) + s/(s-1)$.

Next we prove $\lim_{b\to\infty} g_n(b)/b = n/(s-n)$, $1 \le n < s$. Substitute $a = g_n(b)$ into Eq. (11) and divide it by $b$, we obtain

$$\frac{n(n+1)P_{n+1}(g_n(b),b)}{b} - \frac{n[(n-1)(1+\mu+\nu) + g_n(b) + b + s]P_n(g_n(b),b)}{b}$$
$$= -\frac{s[(n-1)(1+\mu) + g_n(b)]P_{n-1}(g_n(b),b)}{b}.$$

According to $\theta_n(g_n(b),b) = P_n(g_n(b),b)/P_{n-1}(g_n(b),b) = 1$, we obtain

$$\frac{g_n(b)}{b} = \frac{n}{s-n} + \frac{n(n+1)P_{n+1}(g_n(b),b)}{b(n-s)P_n(g_n(b),b)} - \frac{(n-s)(n-1)(1+\mu) + n(n-1)\nu + ns}{b(n-s)}.$$

Letting $b \to \infty$, the boundedness of $P_n$ implies

$$\lim_{b\to\infty} \frac{g_n(b)}{b} = \frac{n}{s-n}.$$

(iii) When $\nu = 0$, we have $\beta = (a+b)/(1+\mu)$, $\omega = s$ and $z_0 = 1/(1+\mu)$. Since $g_1(b) > 0$ increases in $b$, the limit of $g_1(b)$ exists as $b \to 0^+$. Let $g_1(0) = \lim_{b\to 0^+} g_1(b)$. We suppose for contradiction $g_1(0) > 0$ when $\nu = 0$. The monotonicity of $g_1(b)$ in $b$ gives $g_1(b) \geq g_1(0) > 0$, $b > 0$, and the monotonicity of $\theta_1(a,b)$ in $a$ implies

$$1 = \theta_1(g_1(b), b) \geq \theta_1(g_1(0), b), \quad b > 0.$$

$P_n$ tends to Possion distribution when $b \to 0^+$, so $\lim_{b\to 0^+} \theta_1(g_1(0), b) = s > 1$, which obviously contradicts the last equation. Then $g_1(0) = 0$.

Next, we try to verify $\lim_{b\to 0^+} g_1(b)/b = e^{\frac{s}{1+\mu}}/(s-1)$. We have

$$\begin{aligned} P_n(a,b) &= \frac{(\frac{a}{1+\mu})_n}{(\frac{a+b}{1+\mu})_n} \cdot \frac{s^n}{n!} \cdot \frac{{}_1F_1(\frac{a}{1+\mu}+n; \frac{a+b}{1+\mu}+n; -\frac{s}{1+\mu})}{{}_1F_1(\frac{a}{1+\mu}; \frac{a+b}{1+\mu}; s(1-\frac{1}{1+\mu}))} \\ &= \frac{s^n}{n!} \cdot \frac{\Gamma(\frac{a+b}{1+\mu})}{\Gamma(\frac{a}{1+\mu})\Gamma(\frac{b}{1+\mu})} \cdot \frac{\int_0^1 e^{-\frac{st}{1+\mu}} t^{\frac{a}{1+\mu}+n-1}(1-t)^{\frac{b}{1+\mu}-1} dt}{{}_1F_1(\frac{a}{1+\mu}; \frac{a+b}{1+\mu}; s(1-\frac{1}{1+\mu}))} \\ &= \frac{s^n}{n!} \cdot \frac{\Gamma(\frac{a+b}{1+\mu})}{\Gamma(\frac{a}{1+\mu})\Gamma(\frac{b}{1+\mu})} \cdot \frac{\int_0^1 e^{-\frac{st}{1+\mu}} t^{\frac{a}{1+\mu}+n}(1-t)^{\frac{b}{1+\mu}-1} dt + \int_0^1 e^{-\frac{st}{1+\mu}} t^{\frac{a}{1+\mu}+n-1}(1-t)^{\frac{b}{1+\mu}} dt}{{}_1F_1(\frac{a}{1+\mu}; \frac{a+b}{1+\mu}; s(1-\frac{1}{1+\mu}))}. \end{aligned}$$

Let $M(a,b) = {}_1F_1(\frac{a}{1+\mu}; \frac{a+b}{1+\mu}; s(1-\frac{1}{1+\mu}))$. Then

$$P_n(a,b) = \frac{a}{a+b} \cdot \frac{M(a+1+\mu, b)}{M(a,b)} \cdot P_n(a+1+\mu, b) + \frac{b}{a+b} \cdot \frac{M(a, b+1+\mu)}{M(a,b)} \cdot P_n(a, b+1+\mu). \qquad (12)$$

By Eq. (1) with Lebesgue dominated convergence theorem, we obtain

$$\begin{aligned} \lim_{a\to 0^+} P_0(a, 1+\mu) &= \lim_{a\to 0^+} \frac{{}_1F_1\left(\frac{a}{1+\mu}; \frac{a}{1+\mu}+1; -\frac{s}{1+\mu}\right)}{{}_1F_1\left(\frac{a}{1+\mu}; \frac{a}{1+\mu}+1; s\left(1-\frac{1}{1+\mu}\right)\right)} \\ &= \lim_{a\to 0^+} \frac{1 + \sum_{k=1}^{\infty}\left[\frac{\frac{a}{1+\mu}\cdots\left(\frac{a}{1+\mu}+k-1\right)}{\left(\frac{a}{1+\mu}+1\right)\cdots\left(\frac{a}{1+\mu}+k\right)} \frac{(-\frac{s}{1+\mu})^k}{k!}\right]}{1 + \sum_{k=1}^{\infty}\left[\frac{\frac{a}{1+\mu}\cdots\left(\frac{a}{1+\mu}+k-1\right)}{\left(\frac{a}{1+\mu}+1\right)\cdots\left(\frac{a}{1+\mu}+k\right)} \frac{s^k\left(1-\frac{1}{1+\mu}\right)^k}{k!}\right]} = 1. \end{aligned}$$

Similarly, we can obtain

$$\lim_{a\to 0^+} P_n(a, 1+\mu) = 0,\ n \geq 1, \quad \lim_{b\to 0^+} P_n(1+\mu, b) = \frac{s^n}{n!} e^{-s},\ n \geq 0.$$

Let $c > 0$ be a constant. We next consider the limit of $P_n(a,b)$ at $(0,0)$ along the ray $a = cb$. Let $a = cb$, we have

$$\lim_{b\to 0^+} M(cb, b) = \frac{c}{c+1} e^{\frac{\mu s}{1+\mu}} + \frac{1}{c+1}, \quad \lim_{b\to 0^+} M(cb+1+\mu, b) = e^{\frac{\mu s}{1+\mu}}, \quad \lim_{b\to 0^+} M(cb, b+1+\mu) = 1.$$

Taking $a = cb$ into Eq. (12), we find

$$\lim_{b\to 0^+} P_0(cb, b) = \frac{c e^{\frac{\mu s}{1+\mu}} e^{-s} + 1}{c e^{\frac{\mu s}{1+\mu}} + 1}, \quad \lim_{b\to 0^+} P_n(cb, b) = \frac{c e^{\frac{\mu s}{1+\mu}}}{c e^{\frac{\mu s}{1+\mu}} + 1} \cdot \frac{s^n}{n!} \cdot e^{-s},\ n \geq 1.$$

As a result,

$$\lim_{b\to 0^+} \theta_1(cb, b) = \frac{sce^{-\frac{s}{1+\mu}}}{ce^{-\frac{s}{1+\mu}} + 1}.$$

Let $c = e^{\frac{s}{1+\mu}}/(s-1)$, we have

$$\lim_{b\to 0^+} \theta_1(cb, b) = 1 = \theta_n(g_1(b), b),$$

which implies

$$\lim_{b\to 0^+}\frac{g_1(b)}{b}=c=\frac{e^{\frac{s}{1+\mu}}}{s-1}.$$

(iv) Consider $\nu>0$. By the monotonicity of $g_n(b)$, we have $g_n(b_n^*)=\lim_{b\to b_n^*}g_n(b)$ exists, $1\le n<s$. Let $a=(1+\mu)/\nu\cdot b+\omega$, then we have $\alpha=\beta$, and so

$$\theta_n\left(\frac{1+\mu}{\nu}b+\omega,b\right)=\frac{\omega}{n},\quad 1\le n<s.$$

According to the monotonicity of $\theta_n(a,b)$, we obtain

$$0<g_n(b)<\frac{1+\mu}{\nu}b+\omega,\quad 1\le n<\omega$$

and

$$g_n(b)\ge\frac{1+\mu}{\nu}b+\omega,\quad \omega\le n<s.$$

Then we have

$$0\le g_n(b_n^*)=\lim_{b\to b_n^*}g_n(b)\le\frac{1+\mu}{\nu}b_n^*+\omega,\quad 1\le n<\omega$$

and

$$g_n(b_n^*)\ge\frac{1+\mu}{\nu}b_n^*+\omega,\quad \omega\le n<s.$$

By (i), we have $b_1^*=0$ and $b_n^*=0$ if $n\in[\omega,s)$. Thus, we have

$$0\le g_1(0)\le\omega,\ \omega>1,\quad g_1(0)\ge\omega,\ \omega\le 1.$$

Meanwhile, we also have

$$0\le g_n(b_n^*)\le\frac{1+\mu}{\nu}b_n^*+\omega,\quad n\in[2,\omega),\quad g_n(0)\ge\omega,\quad n\in[\omega,s).$$

By (ii), we have $g_1(b)<b/(s-1)+s/(s-1)$, so $g_1(0)\le s/(s-1)$. Thus we have $0\le g_1(0)\le\min\{\omega,s/(s-1)\}$ when $\omega>1$, and $\omega\le g_1(0)\le s/(s-1)$ when $\omega\le 1$. To prove $g_1(0)>0$, we suppose for contradiction that $g_1(0)=0$. Letting $a=g_1(b)$, we have

$$\begin{aligned}&\lim_{b\to 0^+}\theta_1(g_1(b),b)\\ &=\omega\cdot\lim_{b\to 0^+}\left[\frac{\frac{g_1(b)}{1+\mu}}{[g_1(b)+b]z_0+s\nu z_0^2}\cdot\frac{{}_1F_1\left(\frac{g_1(b)}{1+\mu}+1;[g_1(b)+b]z_0+s\nu z_0^2+1;-\omega z_0\right)}{{}_1F_1\left(\frac{g_1(b)}{1+\mu};[g_1(b)+b]z_0+s\nu z_0^2;-\omega z_0\right)}\right]=0.\end{aligned}$$

It contradicts $\theta_1(g_1(b),b)=1$, so $g_1(0)>0$. □

### 4 Proof of Lemma A.2

Recall the definition of $Q_n$ given in Eq. (7). In the following, we rewrite $Q_n$ as $Q_n(a,b)$ to emphasize its dependence on $a$ and $b$. Note that $Q_n(a,b)=0$ defines a contour line of $Q_n$, which is the straight line

$$a=h_n(b)=\frac{nb}{s-n}+\frac{(\mu+\nu)n^2-n(2+\mu+\nu+s\mu)+s(1+\mu)}{s-n},\quad b>\bar{b}_n^*,\tag{13}$$

where

$$\bar{b}_n^* = \max\left\{0, -\frac{(\mu+\nu)n^2 - n(2+\mu+\nu+s\mu) + s(1+\mu)}{n}\right\}.$$

In particular, it is easy to see that $\bar{b}_1^* = 0$ whenever $s > 2$. Let

$$n_1 = \begin{cases} \omega - 1, & \text{there is no interger in the interval } [\omega, s), \\ \omega, & \text{there is at least an interger in the interval } [\omega, s). \end{cases} \tag{14}$$

**Lemma A.2.** Let $s > 2$ and $\mu, \nu \geq 0$ be fixed and let $n_1$ be the constant given in Eq. (14). Let $a = h_n(b)$ be the contour line of $Q_n$ defined above.

(i) Then we have

$$h_n(1+\nu+s\mu) < g_n(1+\nu+s\mu), \quad n \in [1, s),$$
$$g_n(1+\nu+s\mu) < g_{n+1}(1+\nu+s\mu), \quad n \in [1, s-1).$$

(ii) For any $n \in [n_1, s)$, the two curves $a = g_n(b)$ and $a = h_n(b)$ do not intersect. In this case, we have $b_n^* \leq \bar{b}_n^*$ and $g_n(b) > h_n(b)$. Moreover, for any $n \in [n_1, s-1)$, the three curves $a = g_n(b), a = h_n(b)$, and $a = g_{n+1}(b)$ do not intersect. In this case, we have $b_{n+1}^* \leq b_n^* \leq \bar{b}_n^*$ and $g_{n+1}(b) > g_n(b) > h_n(b)$.

(iii) Let $n_1 > 1$. If $a = g_n(b)$ and $a = h_n(b)$ intersect at $b = b_n$ for some $n \in [1, n_1)$, then $a = g_n(b)$, $a = h_n(b)$, and $a = g_{n+1}(b)$ must intersect at the same point. In this case, we have $\bar{b}_n^* \leq b_n^* \leq b_{n+1}^*$ and

$$\begin{aligned} g_{n+1}(b) < g_n(b) < h_n(b), \quad b < b_n, \\ g_{n+1}(b) > g_n(b) > h_n(b), \quad b > b_n, \end{aligned} \tag{15}$$

If $a = g_n(b)$ and $a = h_n(b)$ do not intersect for some $n \in [1, n_1)$, then $a = g_n(b), a = h_n(b)$, and $a = g_{n+1}(b)$ do not intersect. In this case, we have $b_{n+1}^* \leq b_n^* \leq \bar{b}_n^*$ and $g_{n+1}(b) > g_n(b) > h_n(b)$.

*Proof.* (i) Since $b_n^* < 1 + (s-1)\mu < 1 + \nu + s\mu$, we find $b = 1 + \nu + s\mu$ is in the domain of definition of $a = g_n(b)$, where $n \in [1, s)$. For any given $n \in [1, s)$, Taking $b = 1 + \nu + s\mu$ into Eq. (13), we obtain

$$h_n(1+\nu+s\mu) = 1 + \mu + \frac{\mu+\nu}{s-n}n^2.$$

By the definition of $a = h_n(b)$, we have

$$Q_n(h_n(1+\nu+s\mu), 1+\nu+s\mu) = 0.$$

Taking this into Eq. (8), we obtain

$$\begin{aligned} &n(n+1)\Delta P_{n+1}(h_n(1+\nu+s\mu), 1+\nu+s\mu) \\ &= s[(n-1)(1+\mu) + h_n(1+\nu+s\mu)]\Delta P_n(h_n(1+\nu+s\mu), 1+\nu+s\mu). \end{aligned}$$

Then we have either $P_{n+1} \geq P_n \geq P_{n-1}$ at the point $(a, b) = (h_n(1+\nu+s\mu), 1+\nu+s\mu)$, or $P_{n+1} < P_n < P_{n-1}$ at the point $(a, b) = (h_n(1+\nu+s\mu), 1+\nu+s\mu)$. For all $k > n$, we obtain $s - k < s - n$, and so

$$Q_k(h_n(1+\nu+s\mu), 1+\nu+s\mu) = k^2(\mu+\nu) - \frac{n^2(\mu+\nu)(s-k)}{s-n} > 0. \tag{16}$$

Suppose $P_{n+1} \geq P_n \geq P_{n-1}$ at the point $(a, b) = (h_n(1+\nu+s\mu), 1+\nu+s\mu)$. By Eqs. (8) and (16), we can obtain

$$\Delta P_{n+2}(h_n(1+\nu+s\mu), 1+\nu+s\mu) > 0.$$

By using Eqs. (8) and (16) iteratively, we find $\Delta P_k(h_n(1+\nu+s\mu), 1+\nu+s\mu) > 0$ for all $k \geq n+2$. However, Lemma 4.1 implies $\theta_n = P_n/P_{n-1} < s/n \leq 1$ for all $n \geq s$, i.e. $P_n < P_{n-1}$ for all $n \geq s$. This gives a contradiction, so $P_{n+1} < P_n < P_{n-1}$ at the point $(a, b) = (h_n(1+\nu+s\mu), 1+\nu+s\mu)$. Then we have

$$\theta_n(h_n(1+\nu+s\mu), 1+\nu+s\mu) < 1 = \theta_n(g_n(1+\nu+s\mu), 1+\nu+s\mu).$$

The monotonicity of $\theta_n(a, b)$ in $a$ gives

$$g_n(1+\nu+s\mu) > h_n(1+\nu+s\mu), \quad n \in [1, s). \tag{17}$$

Differentiating Eq. (7) with respect to $a$ and $b$, we obtain

$$\frac{\partial Q_n(a,b)}{\partial a} = n - s < 0, \quad \frac{\partial Q_n(a,b)}{\partial b} = n > 0, \quad n \in [1, s).$$

Thus, $Q_n(a, b)$ strictly decreases in $a$ and strictly increases in $b$. By Eq. (17), we have

$$Q_n(g_n(1+\nu+s\mu), 1+\nu+s\mu) < Q_n(h_n(1+\nu+s\mu), 1+\nu+s\mu) = 0.$$

Substituting $b = 1+\nu+s\mu$ and $a = g_n(1+\nu+s\mu)$ into Eq. (6) and combining with above equation, we obtain

$$\theta_{n+1}(g_n(1+\nu+s\mu), 1+\nu+s\mu) < 1 = \theta_{n+1}(g_{n+1}(1+\nu+s\mu), 1+\nu+s\mu).$$

The monotonicity of $\theta_n(a, b)$ in $a$ gives

$$g_n(1+\nu+s\mu) < g_{n+1}(1+\nu+s\mu), \quad n \in [1, s-1). \tag{18}$$

(ii) For $n_1 = \omega - 1$, we obtain there is only one integer in the interval $[\omega - 1, s)$ by the definition of $n_1$. Let $n_2$ be the largest integer in the interval $[1, s)$, then we have $n = n_2$ for all $n \in [\omega - 1, s)$. Taking $n = n_2$ and $a = h_{n_2}(b)$ into Eq. (6), we have

$$n_2(n_2+1)\Big(\theta_{n_2+1}(h_{n_2}(b), b) - 1\Big) = s\Big[(n_2-1)(1+\mu) + h_{n_2}(b)\Big]\Big(1 - \frac{1}{\theta_{n_2}(h_{n_2}(b), b)}\Big). \tag{19}$$

It is obvious that $n_2 + 1 > s$ by the definition of $n_2$, so we have $\theta_{n_2+1}(a, b) < s/(n_2+1) < 1$ for all $a, b > 0$ by Lemma 4.1. This implies $\theta_{n_2+1}(h_{n_2}(b), b) < 1$. Then by Eq. (19), we have

$$\theta_{n_2}(h_{n_2}(b), b) < 1 = \theta_{n_2}(g_{n_2}(b), b).$$

Since Lemma 4.3 proved in the text asserts that $\theta_n(a, b)$ strictly increases with respect to $a$, we have $h_{n_2}(b) < g_{n_2}(b)$. This implies that

$$h_n(b) < g_n(b), \quad n \in [\omega - 1, s).$$

And then we have $b_n^* \leq \bar{b}_n^*$.

Next, we consider $n_1 = \omega$. Inserting Eq. (3) into Eq. (6) gives

$$n^2\theta_n(a+1+\mu, b+\nu) + \frac{s[a + (1+\mu)(n-1)]}{\theta_n(a,b)} = [a + b + s + (n-1)(1+\mu+\nu)]n. \tag{20}$$

Let integer $m \geq 0$. Replacing $n$ by $n+m$ in Eq. (20) gives

$$\begin{aligned}&(n+m)^2\theta_{n+m}(a+1+\mu, b+\nu)\\&= -\frac{s[a+(n+m-1)(1+\mu)]}{\theta_{n+m}(a,b)} + \big[a+b+s+(n+m-1)(1+\mu+\nu)\big](n+m).\end{aligned}$$

Dividing it by $n+m$ and using Eq. (4), we obtain

$$\begin{aligned}&n\theta_n(a+(m+1)(1+\mu), b+(m+1)\nu) + \frac{s[a+(n+m-1)(1+\mu)]}{n\theta_n(a+m(1+\mu), b+m\nu)}\\&= a+b+s+(n+m-1)(1+\mu+\nu).\end{aligned} \tag{21}$$

Differentiating Eq. (13) with respect to $b$, we have $h_n^{'}(b) = n/(s-n)$. Taking $a = h_n(b)$ into Eq. (21) and differentiating Eq. (21) with respect to $b$, we arrive at

$$\begin{aligned}&n\theta_n^{'}(h_n(b)+(m+1)(1+\mu), b+(m+1)\nu)\\&= \frac{s[h_n(b)+(n+m-1)(1+\mu)]\theta_n^{'}(h_n(b)+m(1+\mu), b+m\nu)}{n\theta_n^2(h_n(b)+m(1+\mu), b+m\nu)}\\&\quad+\frac{s}{s-n}\left(1-\frac{1}{\theta_n(h_n(b)+m(1+\mu), b+m\nu)}\right), \quad m\geq 0.\end{aligned} \tag{22}$$

Given $n \in [\omega, s)$. By Lemma $A.1$, we obtain

$$\lim_{b\to\infty}\frac{g_n(b)}{g_{n+1}(b)} = \lim_{b\to\infty}\frac{\frac{g_n(b)}{b}}{\frac{g_{n+1}(b)}{b}} = \frac{n(s-n)-n}{(n+1)(s-n)} < 1,$$

which implies that there is sufficiently large $B_n$ such that when $b > B_n$, we have $g_n(b) < g_{n+1}(b)$. Since $\theta_n(a,b)$ strictly increases with respect to $a$, we obtain

$$\theta_{n+1}(g_n(b), b) < \theta_{n+1}(g_{n+1}(b), b) = 1.$$

Taking $a = g_n(b)$ into Eq. (6) and using the above equation, we have

$$Q_n(g_n(b), b) < 0 = Q_n(h_n(b), b).$$

Since $Q_n(a,b)$ decreases in $a$, we have $h_n(b) < g_n(b)$ for $b > B_n$, which implies that $a = h_n(b)$ and $a = g_n(b)$ do not intersect when $b > B_n$. To prove $h_n(b)$ and $g_n(b)$ have no intersection point, we suppose for contradiction that $a = h_n(b)$ intersects with $a = g_n(b)$. Let $b_n$ be the the largest intersection point, then we have

$$h_n(b) < g_n(b), \quad b > b_n. \tag{23}$$

It is easy to verify $n/(s-n) \geq (1+\mu)/\nu$ as $\omega \leq n$, then taking $b = b_n + m\nu$ into Eq. (13) we have

$$h_n(b_n+m\nu) = h_n(b_n) + \frac{n}{s-n}\cdot m\nu > h_n(b_n) + m(1+\mu), \quad m \geq 1.$$

Combining this with the monotonicity of $\theta_n(a,b)$ in $a$ and Eq. (23), we have

$$\begin{aligned}\theta_n(h_n(b_n)+m(1+\mu), b_n+m\nu) &< \theta_n(h_n(b_n+m\nu), b_n+m\nu)\\&\leq \theta_n(g_n(b_n+m\nu), b_n+m\nu) = 1, \quad m\geq 1.\end{aligned}$$

It then follows from Eq. (22) that

$$\theta_n^{'}(h_n(b_n)+1+\mu, b_n+\nu) = \frac{s[h_n(b_n)+(n-1)(1+\mu)]\theta_n^{'}(h_n(b_n),b_n)}{n^2\theta_n^2(h_n(b_n),b_n)}$$

and

$$\begin{aligned}&\theta_n^{'}(h_n(b_n)+(m+1)(1+\mu), b_n+(m+1)\nu)\\ &< \frac{s[h_n(b_n)+(n+m-1)(1+\mu)]\theta_n^{'}(h_n(b_n)+m(1+\mu),b_n+m\nu)}{n^2\theta_n^2(h_n(b_n)+m(1+\mu),b_n+m\nu)}, \quad m\geq 1.\end{aligned} \tag{24}$$

Suppose $\theta_n^{'}(h_n(b_n),b_n)\leq 0$. Then by using Eq. (24) iteratively, we find that $\theta_n^{'}(h_n(b_n)+m(1+\mu),b_n+m\nu)<0$, where $m\geq 1$. Taking $M>0$ sufficiently large and $\epsilon>0$ such that

$$\Theta = \frac{[h_n(b_n)+M(1+\mu)]+(n-1)(1+\mu)}{s} > 1$$

and

$$\Theta_1 = \max\left\{\theta_n^{'}(a,b_n+M\nu) : a\in\big[h_n(b_n)+M(1+\mu), h_n(b_n)+M(1+\mu)+\epsilon\big]\right\} < 0.$$

Replacing $a$ and $b$ by $a+m(1+\mu)$ and $b_n+M\nu+m\nu$, respectively, in Eq. (24), we obtain

$$\begin{aligned}&\theta_n^{'}(a+m(1+\mu), b_n+M\nu+m\nu)\\ &< \frac{s[a+(n+m-2)(1+\mu)]\theta_n^{'}(a+(m-1)(1+\mu),b_n+M\nu+(m-1)\nu)}{n^2\theta_n^2(a+(m-1)(1+\mu),b_n+M\nu+(m-1)\nu)}.\end{aligned} \tag{25}$$

For any $a\in[h_n(b_n)+M(1+\mu), h_n(b_n)+M(1+\mu)+\epsilon]$, since $\theta_n^{'}(a,b_n+M\nu)<0$, it follows from Eq. (24) that $\theta_n^{'}(a+m(1+\mu),b_n+M\nu+m\nu)<0$. In addition, since $\theta_n<s/n$, it follows from Eq. (25) that

$$\begin{aligned}&\theta_n^{'}(a+m(1+\mu), b_n+M\nu+m\nu)\\ &< [a+(n+m-2)(1+\mu)]\theta_n^{'}(a+(m-1)(1+\mu),b_n+M\nu+(m-1)\nu)\\ &< \Theta^m\theta_n^{'}(a,b_n+M\nu)\\ &< \Theta^m\Theta_1, \quad \forall a\in[h_n(b_n)+M(1+\mu), h_n(b_n)+M(1+\mu)+\epsilon].\end{aligned}$$

By the mean value theorem, there exists $a\in\big(h_n(b_n)+M(1+\mu), h_n(b_n)+M(1+\mu)+\epsilon\big)$ such that

$$\begin{aligned}&\theta_n(h_n(b_n)+M(1+\mu)+\epsilon)-\theta_n(h_n(b_n)+M(1+\mu))\\ &=\epsilon\theta_n^{'}(a+m(1+\mu),b_n+M\nu+m\nu)<\epsilon\Theta^m\Theta_1\end{aligned}$$

Since $\theta_n<s/n$, the left-hand side of the above equation is uniformly bounded for all $m\geq 1$, while the right-hand side tends to negative infinity as $m\to\infty$. This yields a contradiction and hence we have

$$\left.\frac{d\theta_n(h_n(b),b)}{db}\right|_{b=b_n} > 0.$$

As a result, there exists $\delta>0$ such that

$$\theta_n(h_n(b),b)<1, \quad b\in(b_n-\delta,b_n), \quad \theta_n(h_n(b),b)>1, \quad b\in(b_n,b_n+\delta). \tag{26}$$

Since $\theta_n(g_n(b), b) = 1$, by combining Eq. (26) with the monotonicity of $\theta_n(a,b)$ in $b$, we obtain

$$g_n(b) > h_n(b), \quad b \in (b_n - \delta, b_n), \quad g_n(b) < h_n(b), \quad b \in (b_n, b_n + \delta).$$

It also contradicts Eq. (23). Thus $h_n(b)$ and $g_n(b)$ have no intersection point when $\omega \le n < s$. Eq. (18) implies that for $b > 0$ and $\omega \le n < s$, we have

$$h_n(b) < g_n(b).$$

This implies $b_n^* \le \bar{b}_n^*$. Taken together, we conclude that $b_n^* \le \bar{b}_n^*$ and the curves $a = g_n(b)$ and $a = h_n(b)$ do not intersect and satisfy $h_n(b) < g_n(b)$ for any $n \in [n_1, s)$. If the number of integer in the interval $[n_1, s-1)$ is greater than 1, then taking $a = g_n(b)$ into Eq. (6), by the monotonicity of $Q_n(a,b)$ in $a$, we have

$$\theta_{n+1}(g_n(b), b) < 1 = \theta_{n+1}(g_{n+1}(b), b)$$

The monotonicity of $\theta_n(a,b)$ in $a$ indicates $g_n(b) < g_{n+1}(b)$, and we have $h_n(b) < g_n(b) < g_{n+1}(b)$ for $n \in [n_1, s-1)$. This gives $b_{n+1}^* \le b_n^* \le \bar{b}_n^*$, where $n \in [n_1, s-1)$.

(iii) We discuss the case of $1 \le n < n_1$. When $a < (1+\mu)b/\nu + \omega$, we have $\alpha < \beta$ and

$$\theta_n(a + m(1+\mu), b + m\nu) = \frac{\omega}{n} \frac{\int_0^1 e^{-\omega z_0 t} t^{\alpha+m+n-1}(1-t)^{\beta-\alpha-1} dt}{\int_0^1 e^{-\omega z_0 t} t^{\alpha+m+n-2}(1-t)^{\beta-\alpha-1} dt}, \quad m \ge 0.$$

Let

$$I(\lambda) = \frac{s}{n} \cdot \frac{\int_0^1 e^{-st} t^{\lambda+n-1}(1-t)^{\gamma-1} dt}{\int_0^1 e^{-st} t^{\lambda+n-2}(1-t)^{\gamma-1} dt},$$

where $s, \lambda, \gamma > 0$. By the Lemma 4.1 in the reference [4], we obtain that $I(\lambda)$ strictly increases with respect to $\lambda$ for all $s, \gamma > 0$. Let $s = \omega z_0 > 0$, $\gamma = \beta - \alpha > 0$ and $\lambda = \alpha + m > 0$, then $I(\lambda)$ can be rewritten as

$$I(\alpha + m) = \frac{\omega z_0}{n} \cdot \frac{\int_0^1 e^{-\omega z_0 t} t^{\alpha+m+n-1}(1-t)^{\beta-\alpha-1} dt}{\int_0^1 e^{-\omega z_0 t} t^{\alpha+m+n-2}(1-t)^{\beta-\alpha-1} dt} = z_0 \theta_n(a + m(1+\mu), b + m\nu).$$

By the monotonicity of $I(\lambda)$ with respect to $\lambda$, we have

$$\frac{\mathrm{d}\theta_n(a + m(1+\mu), b + m\nu)}{\mathrm{d}m} = \frac{1}{z_0} \cdot \frac{\mathrm{d}I(\alpha+m)}{\mathrm{d}m} = \frac{1}{z_0} \cdot \frac{\mathrm{d}I(\alpha+m)}{\mathrm{d}(\alpha+m)} \cdot \frac{\mathrm{d}(\alpha+m)}{\mathrm{d}m} > 0.$$

Therefore, $\theta_n(a + m(1+\mu), b + m\nu)$ strictly increases with respect to $m$. Then we have

$$\theta_n(a,b) < \theta_n(a + m(1+\mu), b + m\nu), \quad m \ge 1, \tag{27}$$

where $a$ and $b$ satisfy $a < (1+\mu)b/\nu + \omega$. It is obvious that $n_1 \le \omega$ by the definition of $n_1$. The proof of Lemma A.1(iv) gives $g_n(b) < (1+\mu)b/\nu + \omega$ when $n \in [1, n_1)$. If $g_n(b)$ and $h_n(b)$ intersect at $b_n$, then $h_n(b_n) = g_n(b_n) < (1+\mu)b_n/\nu + \omega$. By Eq. (27), we obtain

$$1 = \theta_n(g_n(b_n), b_n) = \theta_n(h_n(b_n), b_n) < \theta_n(h_n(b_n) + m(1+\mu), b_n + m\nu), \quad 1 \le n < n_1.$$

It then follows from Eq. (22) that

$$\theta_n'(h_n(b_n) + 1 + \mu, b_n + \nu) = \frac{s[h_n(b_n) + (n-1)(1+\mu)]\theta_n'(h_n(b_n), b_n)}{n^2 \theta_n^2(h_n(b_n), b_n)}$$

and

$$n\theta_n^{'}(h_n(b_n)+(m+1)(1+\mu),b_n+(m+1)\nu)$$
$$> \frac{s[h_n(b_n)+(n+m-1)(1+\mu)]\theta_n^{'}(h_n(b_n)+m(1+\mu),b_n+m\nu)}{n\theta_n^2(h_n(b_n)+m(1+\mu),b_n+m\nu)}, \quad m \geq 1.$$

Similarly as the proof of (ii), we obtain

$$\frac{d\theta_n(h_n(b),b)}{db}\Big|_{b=b_n} < 0.$$

As a result, there exists $\delta > 0$ such that

$$\theta_n(h_n(b),b) > 1, \quad b \in (b_n-\delta,b_n), \quad \theta_n(h_n(b),b) < 1, \quad b \in (b_n,b_n+\delta). \tag{28}$$

Since $\theta_n(g_n(b),b) = 1$, by combining Eq. (28) with the monotonicity of $\theta_n(a,b)$ in $b$, we obtain

$$g_n(b) < h_n(b), \quad b \in (b_n-\delta,b_n), \quad g_n(b) > h_n(b), \quad b \in (b_n,b_n+\delta). \tag{29}$$

It is then clear that the curve $a = g_n(b)$ can intersect the straight line $a = h_n(b)$ at most once, and $b_n$ is uniquely defined when they do intersect. Then Eq. (29) can be generalized to

$$g_n(b) < h_n(b), \quad b < b_n, \quad g_n(b) > h_n(b), \quad b > b_n. \tag{30}$$

By the definition of $n_1$, we have $n_1+1 < s$, and so the curve $a = g_{n+1}(b)$ exists for all $n \in [1,n_1)$. If $b < b_n$, taking $a = g_n(b)$ into Eq. (6), we obtain

$$\theta_{n+1}(g_n(b),b) > 1 = \theta_{n+1}(g_{n+1}(b),b).$$

Noticing $\theta_n(a,b)$ increases in $a$, so $g_n(b) > g_{n+1}(b)$ when $b > b_n$. A similar argument will verify $g_n(b) < g_{n+1}(b)$ when $b > b_n$. These properties together with Eq. (30) give the ordering of $g_n(b)$, $g_{n+1}(b)$ and $h_n(b)$:

$$g_{n+1}(b) < g_n(b) < h_n(b),\ b < b_n, \quad g_{n+1}(b) > g_n(b) > h_n(b),\ b > b_n, \quad n \in [1,n_1). \tag{31}$$

This indicates that $\bar{b}_n^* \leq b_n^* \leq b_{n+1}^*$, where $n \in [1,n_1)$, and $a = g_n(b)$, $a = h_n(b)$, and $a = g_{n+1}(b)$ intersect at the same point.

If $g_n(b)$ and $h_n(b)$ have no intersection point, then $h_n(b) < g_n(b)$ is obtained by $h_n(1+\nu+s\mu) < g_n(1+\nu+s\mu)$, and so $Q_n(g_n(b),b) < 0$ by the monotonicity of $Q_n(a,b)$ in $a$. By the definition of $n_1$, we have $n_1+1 < s$, and so the curve $a = g_{n+1}(b)$ exists for all $n \in [1,n_1)$. Taking $a = g_n(b)$ into Eq. (6), we obtain

$$\theta_{n+1}(g_n(b),b) < 1 = \theta_{n+1}(g_{n+1}(b),b).$$

The monotonicity of $\theta_n(a,b)$ in $a$ gives $g_n(b) < g_{n+1}(b)$. Thus we have $h_n(b) < g_n(b) < g_{n+1}(b)$ for all $b > 0$ and $1 \leq n < n_1$ if $g_n(b)$ and $h_n(b)$ have no intersection point. This suggests that $b_{n+1}^* \leq b_n^* \leq \bar{b}_n^*$, where $n \in [1,n_1)$, if $g_n(b)$ and $h_n(b)$ have no intersection point. The proof is completed. □

### 5 Proof of Lemma A.3

**Lemma A.3.** Let $s > 2 + 2\nu/(1+\mu)$ and $\mu, \nu \geq 0$ be fixed. Then $n_1 > 1$. Suppose that $a = g_n(b)$ and $a = h_n(b)$ intersect for some $\bar{n} \in [1, n_1)$. Then $a = g_n(b)$ and $a = h_n(b)$ also intersect for any $1 \leq n \leq \bar{n}$. Let $(b_n, a_n)$ be the intersection point of $a = g_n(b)$ and $a = h_n(b)$ in the $b$-$a$ plane for each $1 \leq n \leq \bar{n}$. Then we have

$$a_n < a_{n-1} < \cdots < a_1, \quad b_n < b_{n-1} < \cdots < b_1,$$

and

$$a_1 < \bar{a} = 1 + \left(1 - \frac{2}{s}\right)\mu - \frac{2\nu}{s}, \quad b_1 < \bar{b} = 1 - \left(2 - \frac{2}{s}\right)\nu + \left(s + \frac{2}{s} - 3\right)\mu.$$

*Proof.* Let $(b'_n, a'_n)$ be the intersection point of $a = h_n(b)$ and $a = h_{n+1}(b)$, where $n \geq 1$. By Eq. (13), we obtain

$$b'_n = 1 + \nu + s\mu - \frac{(\mu+\nu)(n+1)(s-n) + (\mu+\nu)ns}{s} < 1 + \nu + s\mu$$

and

$$a'_n = h_n(b'_n) = 1 + \mu - \frac{(\mu+\nu)n(n+1)}{s} < 1 + \mu, \quad n \in [1, s).$$

It is easy to see $a'_n > a'_{n+1}$, so $b'_n > b'_{n+1}$ by the monotonicity of $h_n(b)$ in $b$, where $n \in [1, s)$. Since $s > 2+2\nu/(1+\mu)$, we have

$$a'_1 = 1 + \mu - \frac{2(\mu+\nu)}{s} \geq 1 + \mu - \frac{(\mu+\nu)(1+\mu)}{1+\mu+\nu} = \frac{1+\mu}{1+\mu+\nu} > 0.$$

To complete the proof, we firstly need to verity $b'_n \geq b_n$, where $n \in [1, n_1)$. We suppose for contradiction that there exists $n^* \in [1, n_1)$ such that $b'_{n^*} < b_{n^*}$. Since $h'_n(b) = n/(s-n) < (n+1)/(s-n-1) = h'_{n+1}(b)$, the following relations hold:

$$h_n(b) > h_{n+1}(b),\ b < b'_n, \quad h_n(b) < h_{n+1}(b),\ b > b'_n. \tag{32}$$

By the definition of $b_{n^*}$ and Eq. (32), we have

$$g_{n^*+1}(b_{n^*}) = g_{n^*}(b_{n^*}) = h_{n^*}(b_{n^*}) < h_{n^*+1}(b_{n^*}).$$

Combining this with Eq. (31), we obtain $b_{n^*+1} > b_{n^*}$. Then using Eq. (31) again, for any $b_{n^*} < b < b_{n^*+1}$, we have

$$g_{n^*}(b) < g_{n^*+1}(b), \quad g_{n^*+2}(b) < g_{n^*+1}(b).$$

When $\max\{g_{n^*}(b), g_{n^*+2}(b)\} < a < g_{n^*+1}(b)$, by the monotonicity of $\theta_n(a,b)$ in $a$, we obtain

$$\theta_{n^*}(a,b) > 1, \quad \theta_{n^*+1}(a,b) < 1, \quad \theta_{n^*+2}(a,b) > 1.$$

This gives

$$P_{n^*}(a,b) > P_{n^*-1}(a,b), \quad P_{n^*+1}(a,b) < P_{n^*}(a,b), \quad P_{n^*+2}(a,b) > P_{n^*+1}(a,b),$$

where $b_{n^*} < b < b_{n^*+1}$. Then there exists a nonzero peak at $n = n^*$ and another nonzero peak at $n \geq n^*+2$. However, the steady-state distribution $P_n$ peaks twice at most with possible modes at $n = 0$ and $n = m > 0$, which gives a contrary. Then $b'_n \geq b_n$ for $n \in [1, n_1)$. Since $b'_{n-1} > b'_n$, we have $b'_{n-1} > b'_n \geq b_n$, where $n \in [2, n_1)$. By Eqs. (31) and (32), we obtain

$$g_n(b) > h_n(b) > h_{n-1}(b), \quad b > b'_{n-1}, \quad g_n(b) \leq h_n(b) < h_{n-1}(b), \quad b \leq b_n.$$

Therefore, once $a = g_n(b)$ intersects with $a = h_n(b)$, it intersects with $a = h_{n-1}(b)$ at large $b > b_n$. At the intersection point of $a = g_n(b)$ and $a = h_{n-1}(b)$, Eq. (6) implies that $a = g_{n-1}(b)$ intersects with them there as well. Therefore, $b_{n-1}$ exists with

$$b_n < b_{n-1}, \quad n \in [2, n_1), \tag{33}$$

and the monotonicity of $g_n(b)$ in $b$ gives

$$a_n < a_{n-1}, \quad n \in [2, n_1).$$

Then we conclude that if $a = g_n(b)$ and $a = h_n(b)$ intersect for some $\bar{n} \in [1, n_1)$, then $a = g_n(b)$ and $a = h_n(b)$ also intersect for any $1 \le n \le \bar{n}$, and

$$a_n < a_{n-1} < \cdots < a_1, \quad b_n < b_{n-1} < \cdots < b_1.$$

We next prove $b'_n > b_n$ for all $n \in [1, n_1)$. We suppose there exists $n^* \in [1, n_1)$ such that $b'_{n^*} = b_{n^*}$, then by Eqs. (31) and (32), we have

$$g_{n^*+1}(b'_{n^*}) = g_{n^*+1}(b_{n^*}) = g_{n^*}(b_{n^*}) = h_{n^*}(b_{n^*}) = h_{n^*+1}(b'_{n^*}).$$

Therefore, $b_{n^*+1} = b'_{n^*} = b_{n^*}$, which contradicts Eq. (33). By Eq. (32), we notice that

$$b_1 < b'_1 = 1 + \nu + s\mu - \frac{3(\mu+\nu)(s-1) + \mu + \nu}{s}$$

and

$$a_1 = g_1(b_1) = h_1(b_1) < h_1(b'_1) = 1 + \mu - \frac{2(\mu+\nu)}{s}.$$

The proof is completed. □

### 6 Proof of Lemma A.4

**Lemma A.4.** Let $s > 2$, $\mu, \nu \ge 0$ be fixed, and let $n_2$ be the largest integer in the interval $[1, s)$.

(i) If $2 < s \le 2 + 2\nu/(1+\mu)$, then we have $b^*_n = 0$ and $g_1(b) < g_n(b), b > 0$ for all $n \in [2, s)$.

(ii) If $s > 2 + 2\nu/(1+\mu)$ and $g_n(\bar{b}^*_n) \ge h_n(\bar{b}^*_n)$ for some $n \in [1, s)$, then we have $b^*_n \ge b^*_{n+1} \ge \cdots \ge b^*_{n_2}$ and

$$g_n(b) < g_{n+1}(b) < \cdots < g_{n_2}(b).$$

In particular, if $s > 2 + 2\nu/(1+\mu)$ and $g_1(0) \ge h_1(0) = (s-2)/(s-1)$, then we have $b^*_1 = b^*_2 = \cdots = b^*_{n_2} = 0$ and

$$g_1(b) < g_2(b) < \cdots < g_{n_2}(b), \quad b > 0.$$

*Proof.* (i) Fix $s > 2$ and $\mu, \nu > 0$. Noticing $s \le 2 + 2\nu/(1+\mu)$ only and only if $\omega \le 2$, so we consider $\omega \le 1$ and $1 < \omega \le 2$, respectively. When $\omega \le 1$, we have $[1, s) \subseteq [\omega, s)$. Therefore, by Lemma A.2(ii), we have $a = h_n(b)$ and $a = g_n(b)$ do not intersect for all $n \in [1, s)$, and for all $n \in [1, s-1)$, they satisfy

$$h_n(b) < g_n(b) < g_{n+1}(b).$$

By Lemma A.1(i), we obtain $b_n^* = 0$ for $n \in [1, s)$. Then we have

$$g_1(b) < g_2(b) < \cdots < g_{n_2}(b), \quad b > 0$$

which implies $g_1(b) < g_n(b)$, $b > 0$, for all $n \in [2, s)$.

To proceed, we consider $1 < \omega \leq 2$. Let $a = b(1+\mu)/\nu + \omega$, then we have $\alpha = \beta$, and so

$$\theta_n\left(\frac{1+\mu}{\nu}b + \omega, b\right) = \frac{\omega}{n}.$$

Since $\omega \in (1, 2]$, we find

$$\theta_1\left(\frac{1+\mu}{\nu}b + \omega, b\right) = \omega > 1 = \theta_1(g_1(b), b)$$

and

$$\theta_n\left(\frac{1+\mu}{\nu}b + \omega, b\right) = \frac{\omega}{n} \leq 1 = \theta_n(g_n(b), b), \quad 2 \leq n < s.$$

By the monotonicity of $\theta_n$ in $a$, we have

$$g_1(b) < \frac{1+\mu}{\nu}b + \omega \leq g_n(b), \quad 2 \leq n < s.$$

This indicates that $b_n^* \leq b_1^*$. Lemma A.1(i) suggests that $b_n^* \geq 0$ and $b_1^* = 0$, so $b_n^* = 0$ . Taken together, we conclude that if $2 < s \leq 2 + 2\nu/(1+\mu)$, then $b_n^* = 0$ and $g_1(b) < g_n(b)$, $b > 0$, for all $n \in [2, s)$.

(ii) Consider $s \geq 2 + 2\nu/(1+\mu)$. Then $n_1 \geq \omega - 1 > 1$ and $n_1 \leq \omega < s$. Given $n \in [1, s) = [1, n_1) \cup [n_1, s)$. Firstly, we try to prove when $h_n(\bar{b}_n^*) \leq g_n(\bar{b}_n^*)$, $a = h_n(b)$ and $a = g_n(b)$ do not intersect. According to Lemma A.2(ii), we obtain $a = h_n(b)$ and $a = g_n(b)$ do not intersect if $n \in [n_1, s)$. Then we consider $n \in [1, n_1)$. For any given $n \in [1, n_1)$, we discuss the cases of $h_n(\bar{b}_n^*) < g_n(\bar{b}_n^*)$ and $h_n(\bar{b}_n^*) = g_n(\bar{b}_n^*)$, respectively. Lemma A.2 implies that $a = g_n(b)$ can intersect $a = h_n(b)$ at most once. Therefore, we suppose for contradiction that there exists $b_n > \bar{b}_n^*$ such that $h_n(b_n) = g_n(b_n)$ when $h_n(\bar{b}_n^*) < g_n(\bar{b}_n^*)$. It follows from Lemma A.2 that we have

$$g_n(b) < h_n(b), \quad \bar{b}_n^* < b < b_n.$$

This implies that $h_n(\bar{b}_n^*) \geq g_n(\bar{b}_n^*)$, which contradicts with $h_n(\bar{b}_n^*) < g_n(\bar{b}_n^*)$. Therefore, $a = h_n(b)$ and $a = g_n(b)$ do not intersect when $h_n(\bar{b}_n^*) < g_n(\bar{b}_n^*)$. Consider $h_n(\bar{b}_n^*) = g_n(\bar{b}_n^*)$. By Lemma A.1(iv), we obtain

$$h_n(\bar{b}_n^*) = g_n(\bar{b}_n^*) \leq \frac{1+\mu}{\nu}\bar{b}_n^* + \omega.$$

If $g_n(\bar{b}_n^*) = \bar{b}_n^*(1+\mu)/\nu + \omega$, let $a = g_n(\bar{b}_n^*)$ and $b = \bar{b}_n^*$, then we have $\alpha = \beta$, and so $\theta_n(g_n(\bar{b}_n^*), \bar{b}_n^*) = \omega/n$. Since $n < n_1 \leq \omega$, we have $\theta_n(g_n(\bar{b}_n^*), \bar{b}_n^*) < 1$. This contradicts $\theta_n(g_n(\bar{b}_n^*), \bar{b}_n^*) = 1$, so

$$h_n(\bar{b}_n^*) = g_n(\bar{b}_n^*) < \frac{1+\mu}{\nu}\bar{b}_n^* + \omega.$$

The proof of Lemma A.2(iii) gives $\theta_n(a + m(1+\mu), b + m\nu)$ increases with respect to $m$, so we obtain

$$1 = \theta_n(g_n(\bar{b}_n^*), \bar{b}_n^*) = \theta_n(h_n(\bar{b}_n^*), \bar{b}_n^*) < \theta_n(h_n(\bar{b}_n^*) + m(1+\mu), \bar{b}_n^* + m\nu), \quad m \geq 1. \tag{34}$$

Let $b \to \bar{b}_n^*$, by Eq.(22), we have

$$\begin{aligned} & n\theta_n'(h_n(\bar{b}_n^*) + (m+1)(1+\mu), \bar{b}_n^* + (m+1)\nu) \\ = & \frac{s[h_n(\bar{b}_n^*) + (n+m-1)(1+\mu)]\theta_n'(h_n(\bar{b}_n^*) + m(1+\mu), \bar{b}_n^* + m\nu)}{n\theta_n^2(h_n(\bar{b}_n^*) + m(1+\mu), \bar{b}_n^* + m\nu)} \\ & + \frac{s}{s-n}\left(1 - \frac{1}{\theta_n(h_n(\bar{b}_n^*) + m(1+\mu), \bar{b}_n^* + m\nu)}\right), \quad m \geq 0. \end{aligned}$$

It then follows from Eq. (34) that

$$\theta_n^{'}(h_n(\bar{b}_n^*)+1+\mu,\bar{b}_n^*+\nu)=\frac{s[h_n(\bar{b}_n^*)+(n-1)(1+\mu)]\theta_n^{'}(h_n(\bar{b}_n^*),\bar{b}_n^*)}{n^2\theta_n^2(h_n(\bar{b}_n^*),\bar{b}_n^*)}$$

and

$$\begin{aligned}&n\theta_n^{'}(h_n(\bar{b}_n^*)+(m+1)(1+\mu),\bar{b}_n^*+(m+1)\nu)\\&>\frac{s[h_n(\bar{b}_n^*)+(n+m-1)(1+\mu)]\theta_n^{'}(h_n(\bar{b}_n^*)+m(1+\mu),\bar{b}_n^*+m\nu)}{n\theta_n^2(h_n(\bar{b}_n^*)+m(1+\mu),\bar{b}_n^*+m\nu)},\quad m\geq 1.\end{aligned}$$

Similarly as the proof of Lemma A.2(ii), we obtain

$$\theta_n^{'}(h_n(\bar{b}_n^*),\bar{b}_n^*)<0.$$

As a result, there exists $\delta>0$ such that

$$\theta_n(h_n(b),b)<1,\quad b\in(\bar{b}_n^*,\bar{b}_n^*+\delta).$$

Since $\theta_n(g_n(b),b)=1$ and $\theta_n(a,b)$ increases strictly with respect to $a$, we obtain

$$h_n(b)<g_n(b),\quad \bar{b}_n^*<b<\bar{b}_n^*+\delta. \tag{35}$$

If $a=h_n(b)$ and intersect with $a=g_n(b)$ at $b=b_n>\bar{b}_n^*$, then we have

$$h_n(b)>g_n(b),\quad \bar{b}_n^*<b<b_n.$$

This contradicts Eq. (35), so $a=h_n(b)$ and $a=g_n(b)$ do not intersect. Taken together, we conclude that $a=h_n(b)$ and $a=g_n(b)$ do not intersect when $g_n(\bar{b}_n^*)\geq h_n(\bar{b}_n^*)$ for any given $n\in[1,s)$. It then follows from Lemma A.2 that

$$h_n(b)<g_n(b)<g_{n+1}(b),\quad n\in[1,s-1). \tag{36}$$

when $h_n(\bar{b}_n^*)\geq g_n(\bar{b}_n^*)$.

Second, we prove if $h_n(\bar{b}_n^*)\leq g_n(\bar{b}_n^*)$, then $h_{n+1}(\bar{b}_{n+1}^*)\leq g_{n+1}(\bar{b}_{n+1}^*)$, where $n\in[1,s-1)$. We suppose for contradiction that $h_{n+1}(\bar{b}_{n+1}^*)>g_{n+1}(\bar{b}_{n+1}^*)$. By $h_{n+1}(1+\nu+s\mu)<g_{n+1}(1+\nu+s\mu)$ given in Lemma A.2(i), we find $a=h_n(b)$ intersects with $a=g_n(b)$ at $b\in(\bar{b}_n^*,1+\nu+s\mu)$. By Lemma A.2, we have $a=h_{n+1}(b)$ and $a=g_{n+1}(b)$ intersect exactly once. However, Lemma $A.3$ indicates that, once $a=h_{n+1}(b)$ intersects with $a=g_{n+1}(b)$, we have $a=g_n(b)$ and $h_n(b)$ intersect, which gives a contradiction . Therefore, if $h_n(\bar{b}_n^*)\leq g_n(\bar{b}_n^*)$, then $h_{n+1}(\bar{b}_{n+1}^*)\leq g_{n+1}(\bar{b}_{n+1}^*)$. Following the similar discussion in Eq. (36), we obtain $h_{n+1}(b)<g_{n+1}(b)<g_{n+2}(b)$. Repeating this process, we can obtain when $h_n(\bar{b}_n^*)\leq g_n(\bar{b}_n^*)$,

$$g_n(b)<g_{n+1}(b)<\cdots<g_{n_2}(b),\quad n\in[1,s).$$

This indicates that $b_n^*\geq b_{n+1}^*\geq\cdots\geq b_{n_2}^*$. When $n=1$, we have $b_1^*=\bar{b}_1^*=0$ by Lemma $A.1$(i) and Eq. (13). Therefore, if $g_1(0)\geq h_1(0)=(s-2)/(s-1)$, we have for all $b>0$

$$g_1(b)<g_2(b)<\cdots<g_{n_2}(b),\quad n\in[1,s)$$

The proof is completed. □
